# Divergence of the individual repeats in the leucine-rich repeat domains of human Toll-like receptors explain their diversity and functional adaptations

**DOI:** 10.1101/2024.09.30.615863

**Authors:** Abraham Takkouche, Keita Ichii, Xinru Qiu, Lukasz Jaroszewski, Adam Godzik

## Abstract

Toll-like receptors (TLRs) are best known pattern recognition receptors of innate immunity, detecting a broad range of pathogen-associated molecular patterns (PAMPs), and endogenous danger associated molecular patterns (DAMPs), initiating inflammatory and antimicrobial responses. TLRs contain two specialized domains, a signaling domain (TIR), and a receptor domain composed of tandem repeats of short 20-30 amino acid segments called Leucine-rich Repeats (LRRs). LRR domains, often paired with other domains, are widespread in all kingdoms of life, invariably forming highly similar, solenoid-like three dimensional structures. Despite this structural conservation, LRR domains overall, and receptor domains of TLRs in particular, exhibit remarkable diversity in binding specificity, recognizing diverse ligands such as lipoproteins, nucleic acids, polysaccharides, other proteins and protein complexes. To understand how this conserved scaffold can accommodate such binding diversity, we performed an in-depth analysis of sequential and structural conservation of individual repeats within the LRR domain of each of the ten human TLRs. We demonstrate that the small variations in repeat lengths and local sequence patterns lead to subtle, but critical structural adaptations, such as changes in local curvature, emergence of loops, cavities and interaction interfaces, each contributing to recognition of specific ligands and formation of the receptor complexes. TLR polymorphisms in human populations can further fine-tune the specificity and strength of ligand recognition, influencing how individuals respond to different pathogens and cell damage causing diseases. We show that in most cases the interfaces with the ligands show high level of polymorphism, suggesting a potential for diverse immune responses to infections among human populations while interfaces with other proteins in the receptor complex are more conserved, pointing to the importance of conserving the overall structure of the signaling pathways. By studying how divergence in LRR repeats affects TLRs structure and function, we provide deeper insights into TLR recognition mechanisms, and a better understanding of the mechanism of evolutionary adaptability of immune recognition systems, both along the evolution of vertebrates, and across the human population.

**Significance statement:** Human Toll-like receptors (hTLRs) play a key role in the innate immune response, recognizing diverse danger molecular patterns through their receptor domains, that consist of tandem repeats of a structural unit called a leucine rich repat (LRR). They are also an example of functionally diverse paralogous family where, despite the overall sequence and structure similarity, each member develops its specific function. Our study reveals that subtle modifications of dividual repeats, without disrupting the overall structure of the receptor domain, form secondary patterns defining the functional specificity of each TLR, enabling them to recognize and respond to a broad array of pathogen and disease associated molecular patterns. These findings offer a new perspective on how sequence variability within conserved protein domains can drive functional evolution and adaptation, with implications both for understanding immune receptor functional adaptation and their fast evolution as well as more general problem of functional diversification in paralogous families.

## Introduction

Activation of innate immunity is the first, critical step in a defense against pathogens and tissue damage. This is achieved by a multitude of Pattern Recognition Receptors (PRRs) (1), specialized proteins that recognize molecular signatures of pathogens, Pathogen-Associated Molecular Patterns (PAMPs), and other conserved molecular patterns associated with danger to the cell, Damage-Associated Molecular Patterns (DAMPs). Upon recognizing such patterns, the receptors become activated and initiate downstream signaling cascades that turn on appropriate sections of the immune network, resulting in the release of specific cytokines or activating autophagy, cell death networks, and phagocytosis (2-5). One of the best studied groups of innate immunity receptors are Toll-like receptors (TLRs), named after a Drosophila Toll protein. All TLRs consist of two specialized domains: the N-terminal leucine-rich repeat (LRR) domain that recognizes specific PAMPs and DAMPs and a C-terminal TIR (toll-IL-1 receptor) domain that passes the signal down through binding to the intra-cellular receptors, connected by a single transmembrane helix.

The ligand binding N-terminal domains of TLRs are composed of multiple repeats of a sequence and structure motif called a **L**eucine **R**ich **R**epeat (LRR). They belong to the broad class of tandem repeat proteins and specifically to the α/β solenoids of the LRR RI-like subfamily (6), members of class III of tandem repeat proteins (7). LRR domains of TLRs directly or indirectly recognize ligands associated with different pathogens, such as bacterial Lipopolysaccharides (LPS), lipoproteins, zymosan, flagella, single and double stranded RNA and other PAMPs, as well as Heat shock proteins (HSPs), extracellular matrix components, nucleic acids, and other DAMPs from damaged or dying cells (8). LRR domains are present in many proteins with other architectures, including over 300 diverse proteins in the human proteome. Toll-like receptors emerged at the base on the Cnidaria/Bilateria but immune receptors with LRR and TIR domains evolved independently in plants (R-proteins). Plants and invertebrates utilize large numbers of receptors, with plants employing thousands of R-proteins and some invertebrates employing hundreds of TLRs, most of which are specifically tailored to recognize a single pathogen (9). In contrast, mammals possess only 10 to 15 TLR receptors capable of recognizing a broad spectrum of pathogens. Many of these receptors can bind to and be activated by a diverse set of both endo- (10) and exo-genous (11) ligands.

Analyses and comparisons between tandem repeat proteins are challenging due to their periodic structure because of their translational symmetry, both sequence and structural alignments remain similar even when shifted by multiples of the repeat length (12). This periodicity can obscure true evolutionary relationships, as the repeating pattern may dominate the similarity signal. This affects phylogeny analysis and leads to many misnamed proteins, where, for instance, apparent orthologs of vertebrate TLRs are identified in invertebrates while more careful phylogenetic analysis shows that they are a part of species-specific expansion (13). In this contribution we focus on another, rarely discussed feature of tandem repeat proteins – the divergence of the individual repeats within specific proteins and its consequences for their functions. In tandem repeat proteins, after their initial emergence from identical copies of a single repeat, each repeat evolves under individual, function-driven constraints. Depending on these constraints, some LRR domains are very regular, with all repeats having the same length and conserved sequence pattern, while in others, individual repeats can be highly divergent. In Fig. 1 we present a comparison between an archetypal, regular LRR protein, the ribonuclease inhibitors (RNH1) and one of the human TLR proteins, TLR5.

**Figure 1.**
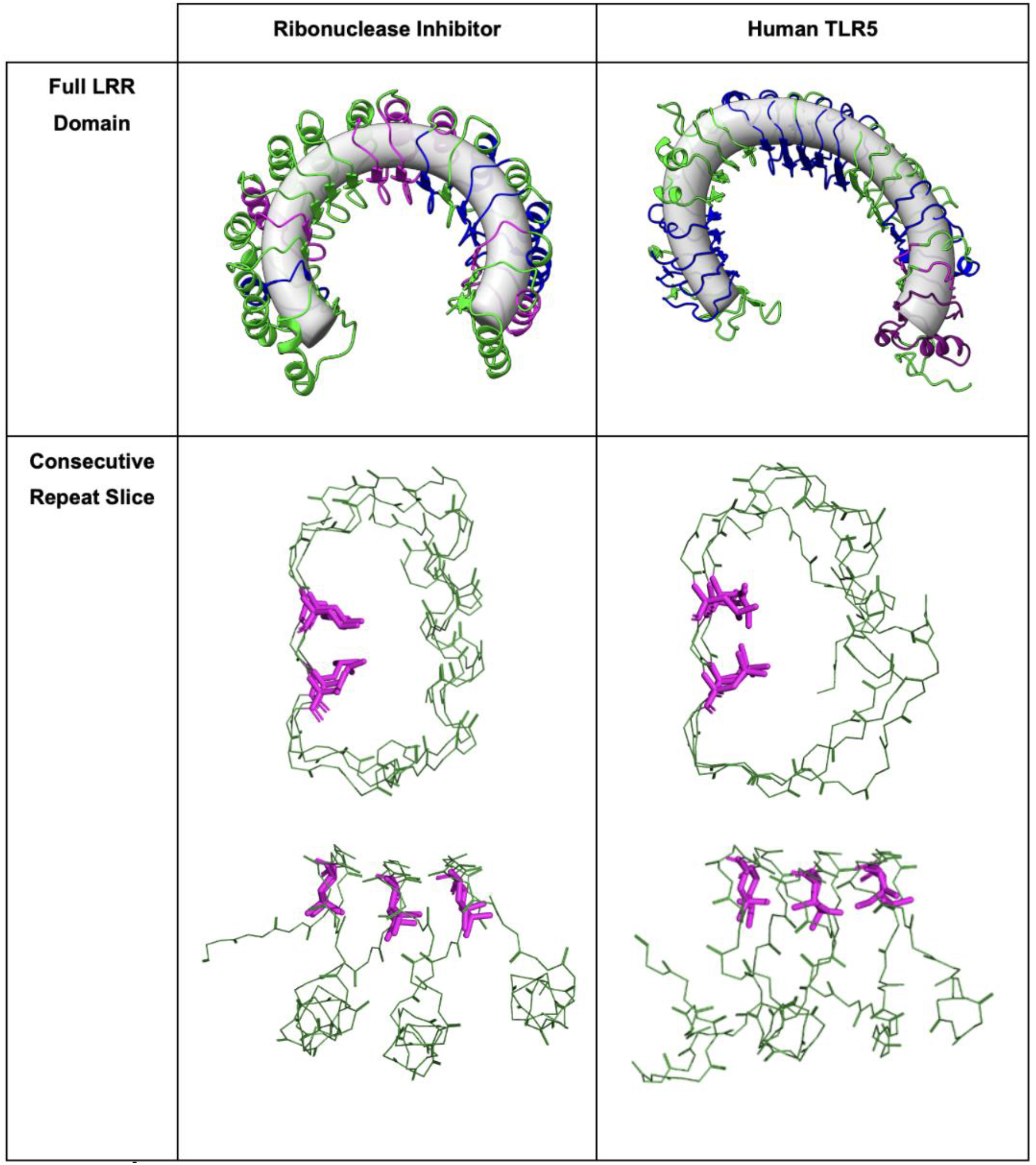
Examples of the regular and irregular LRR domains. Top left: A regular LRR domain, the porcine ribonuclease inhibitor (PDB:1DFJ), the first LRR protein with an experimentally solved structure (14). The LRR domain of the RNH1 structure is superimposed over a 3D model of a regular torus. 8 repeats out of 17 repeats are recognized by InterPro/Pfam HMMs, with colors showing different subtypes. Lower left: A structural representation of three consecutive repeats, spanning positions 86 to 141 is shown in the bottom panel, with the leucines in the LxL motifs highlighted in magenta. Top right: An example of highly irregular LRR domain of hTLR5 (PDB:3J0A) superimposed with an ellipsoid torus. 7 out of 21 repeats are recognized by InterPro/Pfam HMMs in hTLR5. Lower right: A structural representation of three consecutive repeats, spanning positions 498 to 560 is shown in the bottom panel, with the leucines in the LxL motifs highlighted in magenta.

**Figure 2.**
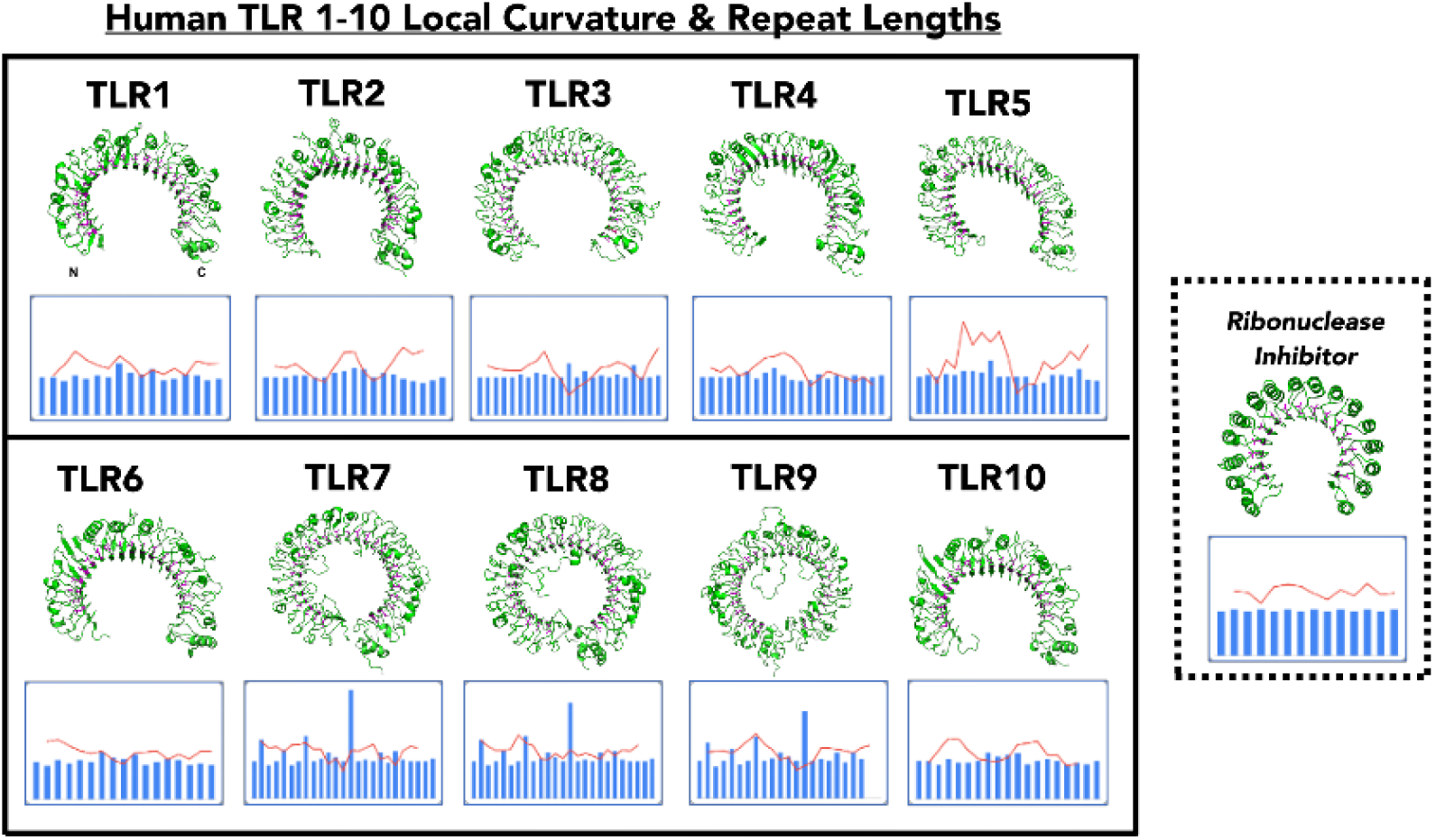
Thumbnail figures of the three-dimensional structures of human TLR 1-10, illustrating the distortions from the ideal solenoid structure (RNH1), right panel. Local curvature moving averages and repeat lengths (see the Methods section) are rendered as red curve on scale from 0 to 0.1, and blue bars on scale from 0 to 75, respectively are shown below each structure. Full data is available in dataset 1 of the supplemental materials. AlphaFold2 models of the structures of hTLR5, hTLR6, hTLR9 and hTLR10 are used here, as no experimental structures were available as of this writing.

Several algorithms (15), including some developed in our group (16), focus on identifying individual repeats in tandem repeat proteins, enabling the analysis of their divergence. As shown in Figure 1, the human TLR5 (hTLR5) Pfam HMMs (15) two different HMM models from the 16 available in the Pfam database recognized only 7 (out of 21) individual LRRs, illustrating repeat divergence, as observed. Even in the more regular RNH1, Pfam recognized only 6 out of the 14 LRRs (17). In comparison, SMART HMMs performed slightly better, recognizing 9 repeats in TLR5 and 13 in RNH1. Human analysis and algorithms analyzing the 3D structure provide the ultimate answer to recognizing all the repeats, and some of the more recent, AI based algorithms can match them in their ability to identify all the repeats based on the sequence alone (18, 19).

The structural diversity of individual repeats can be subtle and is often not captured by the protein structure comparisons that rely on global similarity measures such as RMSD or TM-score (20), as they align most LRR proteins to each other with RMSDs below 3Å and a high TM-score, indicative of “strong” structural similarity. However, these apparently minor structural differences are critical for the functional diversity of the TLR LRR domains. The focus of this manuscript is on the description and analysis of differences between individual repeats in tandem repeat proteins at the levels of sequence, structure, and function. In particular, we examine the correlation between sequence divergence, the local curvature of the concave side of the solenoid, and the presence of functional motifs— such as extra-long loops, binding sites, or cavities—often referred to as “embellishments” of the core fold (21). We also investigate how these divergences correspond to sequence polymorphisms observed in the human population. It is well established that humans (and other animals) exhibit substantial variation in immune responses to the same pathogens, and that genes involved in immunity are under strong positive selection and display high level of polymorphisms within populations (22). As genome sequencing expands to include larger and more geographically and ethnically diverse cohorts, our understanding of human genetic variation continues to grow. In this study, we show that the frequency of mutations and other genomic divergence in human populations correlates with the structural diversity of individual repeats and functionally important positions in human TLRs. This supports epidemiological observations suggesting that such diversity contributes to adaptation to distinct pathogens across different environments. Mapping and analyzing these differences are essential for understanding the functional divergence of TLR immune receptors and their implications for human health.

Consensus patterns for different LRR protein subtypes (6), focused primarily on the namesake conserved leucine pattern. For the RI-like subtype of LRR proteins such as TLRs, the consensus motif has been defined as LxxLxLxxN/CxL, with some definitions extending towards the C-terminus by incorporating a secondary, minor, pattern, oxxLxxoL (6). Occasional presence of other amino acids in classical leucines’ positions was noted, but such breaks in the pattern were usually described as random divergence. Here, we argue that changes in this pattern contribute to the nuances of the local structure of the LRR domains, and the repeats breaking the pattern are critical to the specific functions of the individual TLRs and other LRR proteins.

<u>Here, we focus on the following features of individual repeats</u>

– **Starting points and length of individual repeats**.
– **Conservation of the consensus sequence pattern**.
– **The local curvature of the inside of the solenoid**.
– **Presence of extra-long loops** extending beyond the main solenoid body
– **Positions involved in ligand binding and complex stabilization**.
– **Levels of polymorphisms in human population** in individual repeats or around specific functional features.

### Starting point and length of individual repeats

Selecting a starting point in the periodic function is arbitrary, so we decided to define the start of each repeat at the first amino acid in the LxL pattern and defined the length as a distance to the equivalent residue in the following repeat. The central “LxL” pattern, including the divergent patterns, was identified for each structure using a combination of a regex-based pattern search, the solenoid structure repeat searching algorithm ConSole (16) and new algorithms based on unsupervised clustering of pLM embeddings (18, 19) and confirmed through manual analysis with a molecular visualization software, PyMol. The entire sequence of the LRR domains of all the human TLRs was parsed into individual repeats based on these positions.

### Conservation of the consensus sequence pattern

Number of conserved positions out of 5 in the LxxLxLxxN/CxL pattern defining the RI-like subtype of LRR proteins (6).

### The local curvature of the inside of the solenoid

We calculated the local curvature of the LRR domain solenoids by placing a circle going through Cα atoms of the residues defining a start of three consecutive repeats. We have also tested using Cβ atoms or taking ± 2 repeats or fitting a circle to 5 consecutive repeats, all leading to comparable results (data not shown). A circumradius, or its inverse (Menger curvature), was used to define the local curvature (23). To smooth fluctuations and reduce noise in visualizing local curvature across the length of the LRR domain, we applied a moving average with an interval of three. The moving average was computed by taking the mean of each data point with its two neighboring points. The 3-dimensional coordinates of the structure were obtained from the PDB database (24). For predicted models, coordinates were either retrieved from the AlphaFold database (23) or generated locally using the publicly available AlphaFold2 software (24).

To validate our method for measuring local curvature, we examined the structure of ribonuclease inhibitor (RNH1) (PDB:1DFJ) as a baseline, regarded as a classical RI-like LRR in terms of both sequence and structure (14, 25). As expected, the local radius was consistent across the entire length of the protein. Additionally, the consensus sequence pattern was conserved in all repeats, and their lengths were systematically alternating between 30 and 31.

There are no experimental structures for human TLR6, TLR9 and TLR10. To have a complete set of structures for the analysis, we used AlphaFold2 (AF2) predicted structures of these proteins as downloaded from the UniProt databae. To test if the quality of these models is sufficient to reproduce local structural features, we performed a series of comparisons between the local curvature of experimental models versus AI-predicted structures for the several RI-like subtype of LRR domains with experimentally determined structures which showed very little divergence. In a comparison of the experimental and AlphaFold2 models of TLR1 for instance, the two local curvature datasets were found to have no significant difference with a p-value of 0.94. The data for these analyses are provided in Dataset 2 of the Supplemental Materials.

### Presence of extra-long loops

extending beyond the main solenoid body was deduced from the repeat lengths and verified by the manual analysis of individual TLR protein models.

### Positions involved in ligand binding and complex stabilization

Residues involved in ligand interactions and protein-protein complexes have been identified from the analysis of experimental structures downloaded from the PDB (if available) or from AlphaFold3 models of complexes. A newer versions of the AlphaFold algorithm was used to model protein complexes, as it can handle larger (up to 5,000 amino acids) systems. The latter cases are explicitly identified in the text as AF3 predictions. Residues with at least one heavy atom within a 4.5Å distance of any heavy atom in the ligand or in the other chain in the complex were identified as involved in the binding interface.

### Effects of polymorphism in human population

Aggregated data on the genomic diversity in human population was downloaded from gnomAD database (26). Version v4, based on the GRCh38 human genome assembly, contained data from 730,947 exome sequences and 76,215 whole-genome sequences which were used in this analysis. Missense mutations were selected from the data and assigned to corresponding residues in experimental structures (or AF3 models) of complexes of TLRs with their ligands.

The average mutation frequency for a residue was defined as the mean frequency for 15 residues from the same chain with the shortest spatial distance to this residue. Then, interfaces between proteins in complexes or proteins and ligands were defined as sets of residues within 4.5 Å from any heavy atom in the interacting molecules. Average mutation frequencies in residues involved in each interaction interface (represented as orange dots in figures 3-7) were then compared with average mutation frequencies in residues outside this interface (represented as grey dots in figures 3-7) using a two-tailed t-test to assess the significance of the difference between these two sets. All distances between protein residues and ligands were defined as the minimum distance between their heavy atoms.

**Figure 3.**
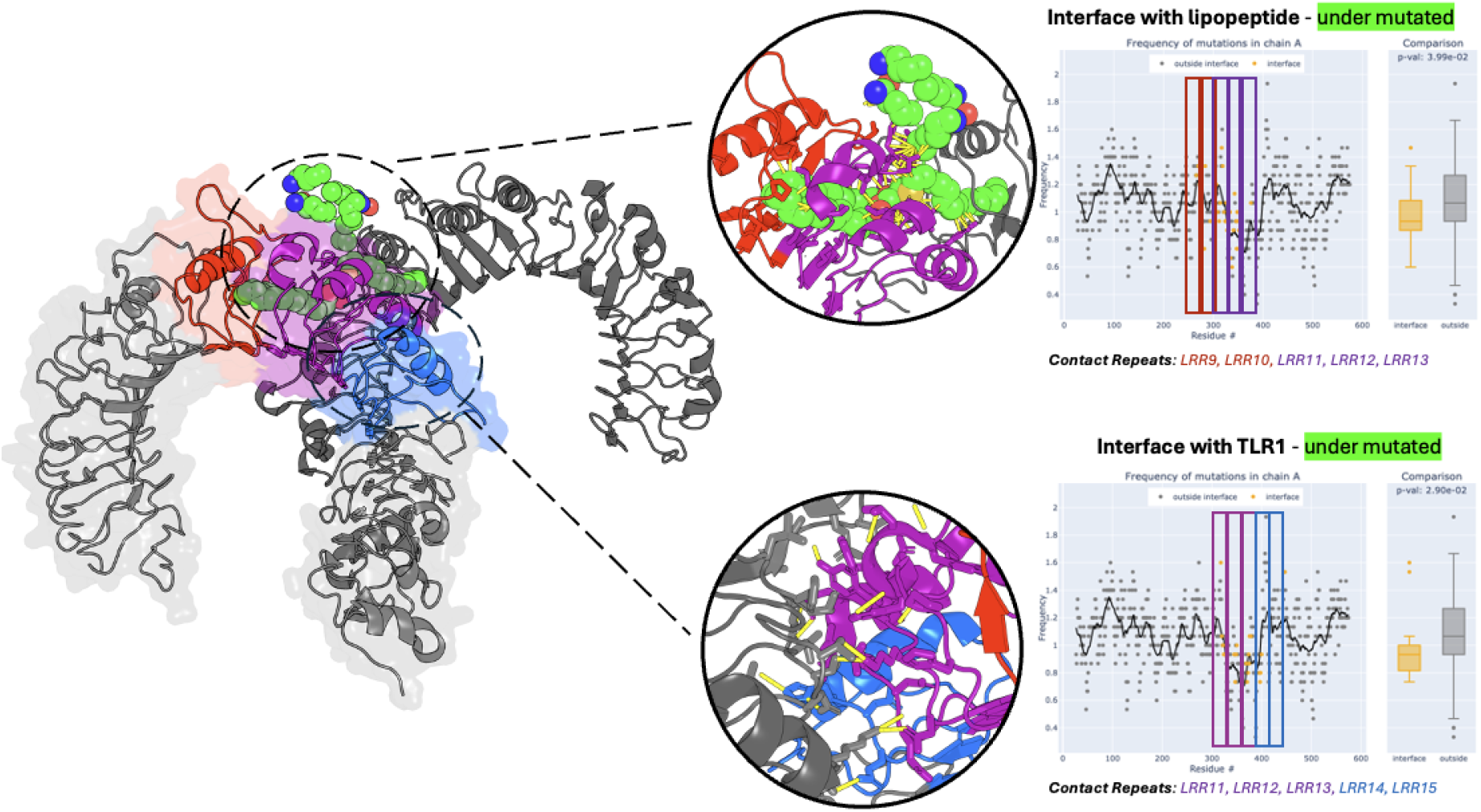
hTLR1-hTLR2 heterodimer (PDB: 2Z7X (30)) with the insets on the right showing the distribution of mutations along the length of hTL2, highlighting key interaction sites with the ligand and complex interfaces. The LRRs in hTLR2 that interact with hTLR1 are shown in blue, while those interacting with the lipopeptide ligand are shown in red. LRRs involved in both interactions are highlighted in purple. These contact repeats are labeled below the mutation frequency plot and boxed within it. Close-up views of these interactions are circled for clarity. In human TLR2, both the dimerization interface and the lipopeptide-binding site show reduced frequency of mutation. LRR12 and LRR9-10, are the longest and most divergent repeats in TLR2. Loops between them bind the lipoprotein head and shape the opening of the ligand-binding cavity. These repeats introduce a kink in the inner surface of the solenoid and form the cavity that accommodates the hydrophobic portion of the ligand. The divergent repeats spanning LRRs 9 to 12 interact with the LPS tail with their high level of polymorphism combined may facilitate recognition of structurally diverse forms of LPS.

**Figure 4.**
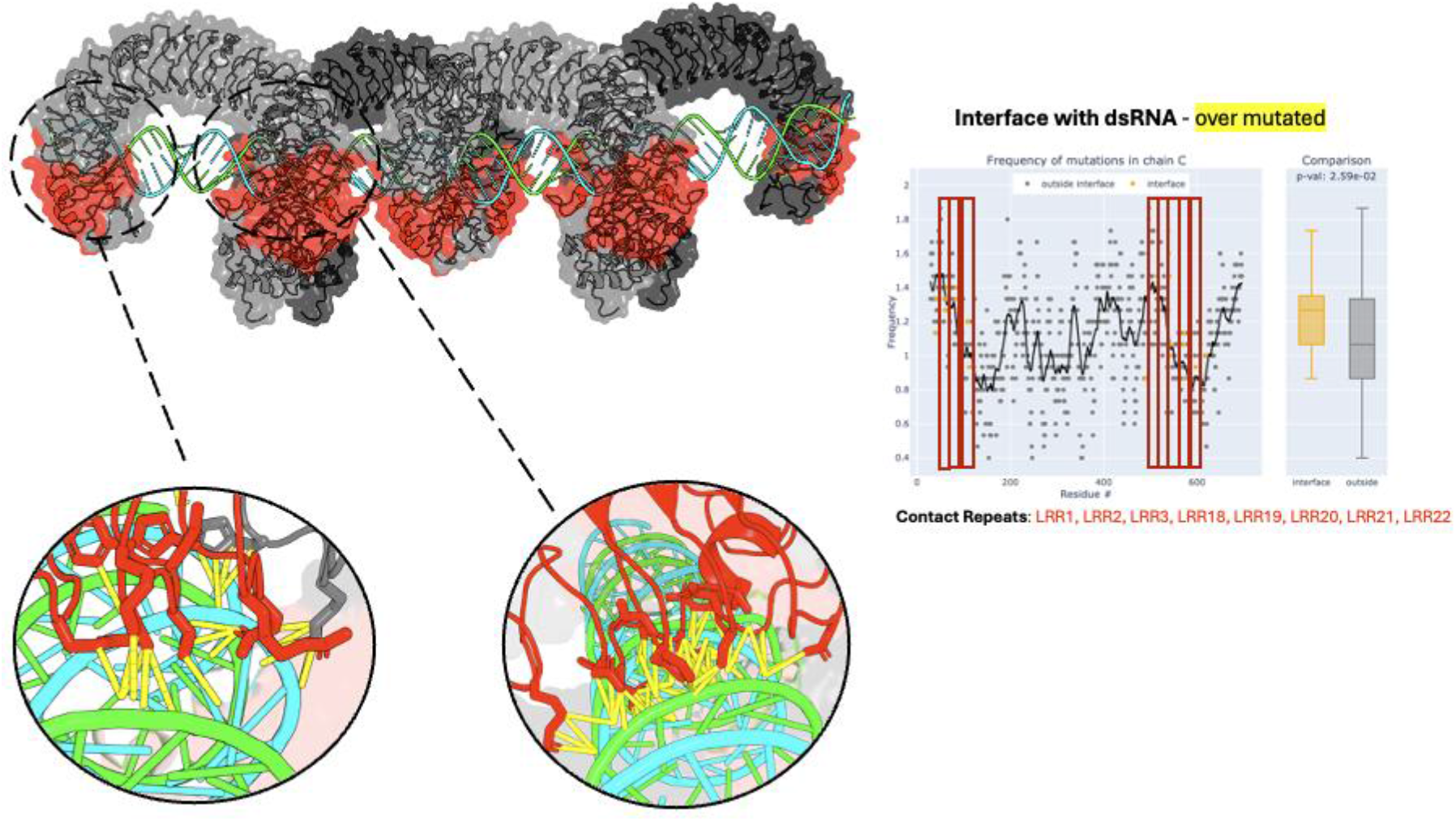
Structural representation of the hTLR3 LRR domains in complex with RNA (PDB: 7WV3 (32)) with the distribution of mutations plotted along the length of hTLR3. The dsRNA ligand interface is highlighted in red. These contact repeats are labeled below the mutation frequency plots and identified by red boxes. RNA-binding regions in all four chains are over mutated; however, only chain A is shown in this representation. Interactions between hTLR3 chains were not found to be significant. Close-up views of these interactions are circled for clarity. Most of the interaction driving residues are located on LRR20, the longest and most divergent repeat in hTLR3.

**Figure 5.**
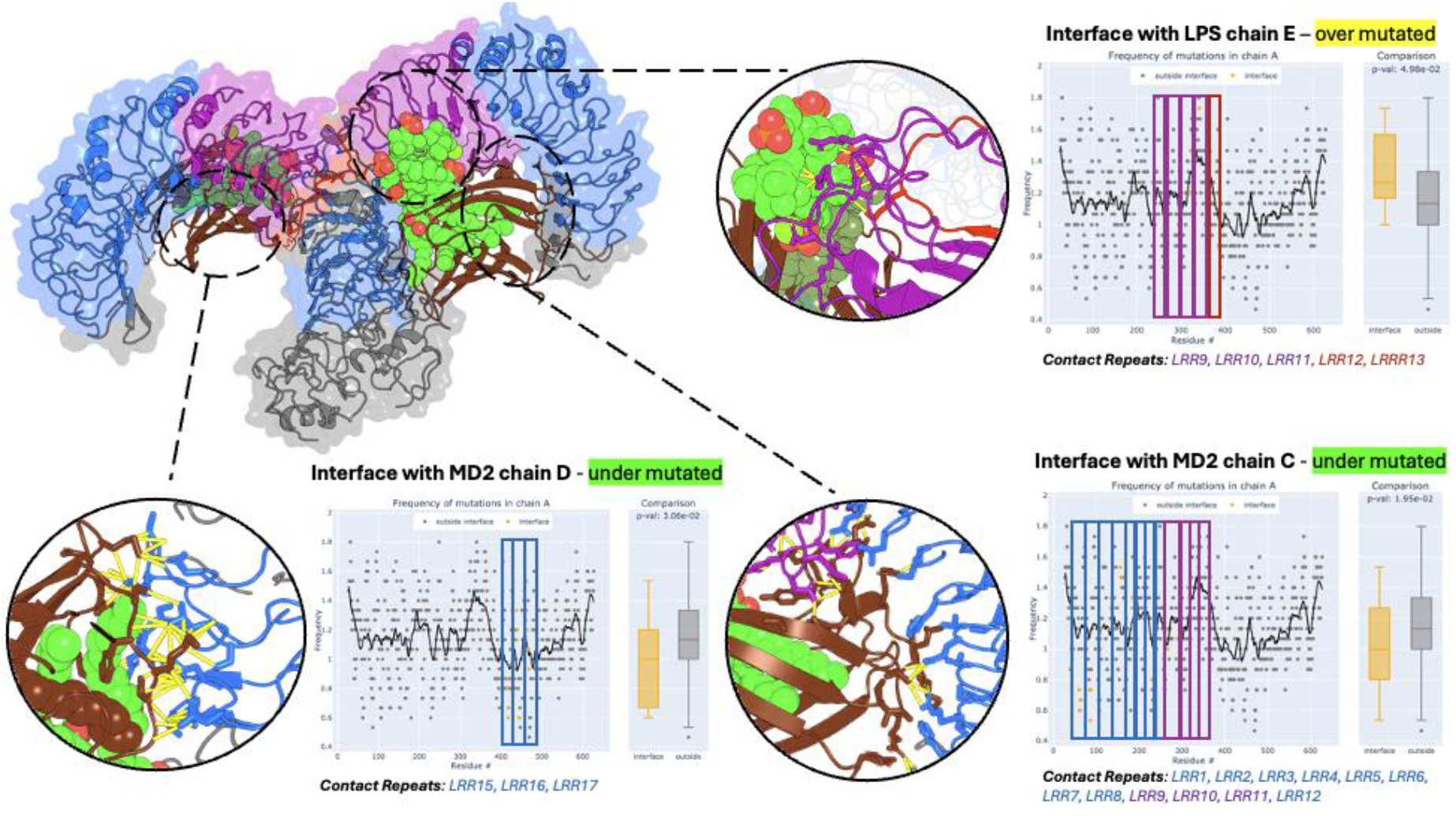
Structural representation of the hTLR4-MD2-LPS complex (PDB:3FXI (36)) with the distribution of mutations plotted along the length of hTLR4. The LRRs in hTLR4 that interact with MD2 are shown in blue, while those interacting with the LPS head are shown in red. LRRs involved in both interactions are highlighted in purple. These contact repeats are labeled below the mutation frequency plot and boxed within it. Close-up views of these interactions are circled for clarity. Mutation frequency of the interactions between hTLR4 chains were not found to be significantly different from the rest of the hTLR4. The most structurally divergent repeats in hTLR4 (LRR9-LRR13) form functional loops that create the dimerization interface for interactions with the LPS head (top right close-up). The LRR repeat with the loop stabilizing TLR4/MD2 interface (LRR9) and one with the loop providing the “lid” to the LPS binding cavity in MD2 (LRR17) are the longest and the most sequentially divergent of all the repeats, respectively.

**Figure 6.**
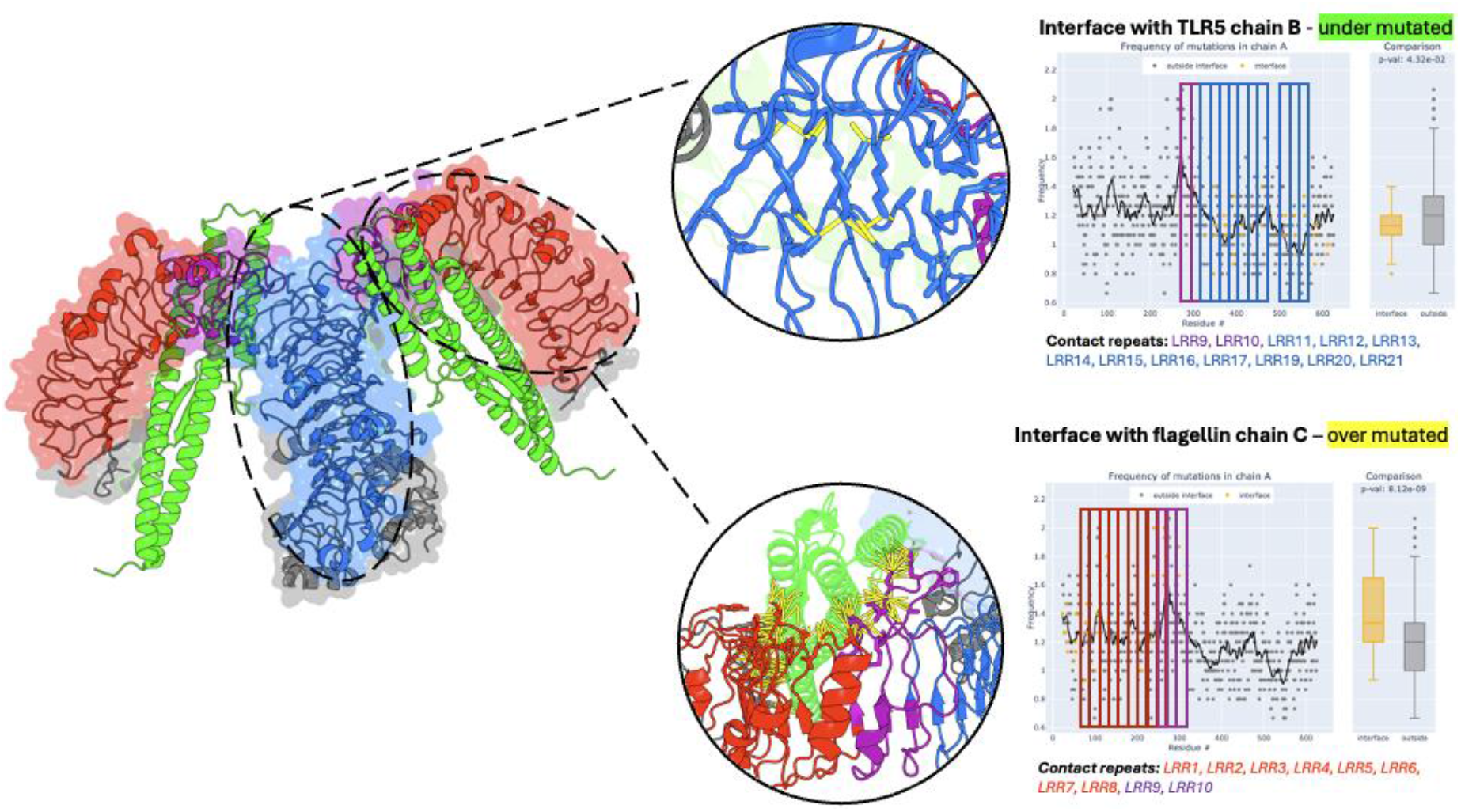
Structural representation of the hTLR5-flagellin complex. As the only available experimental structure of a TLR5/flagellin complex with resolution sufficient to study local structure details comes from zebrafish (37), and human and zebrafish TLR5 are not strongly similar (35% seq id), an (AF3) model of the analogous complex of human TLR5 is shown here. Individual repeats involved in interfaces between the two hTLR5 chains are shown in blue, while those interacting with bacterial flagellin are shown in red. LRRs involved in both interactions are highlighted in purple. These contact repeats are labeled below the mutation frequency plot and boxed within it. Close-up views of these interactions are shown in more details for clarity. Interaction interfaces with flagellin and the critical loop forming TLR5 homodimer interface upon binding to flagellin, are over- and under mutated, respectively.

**Figure 7.**
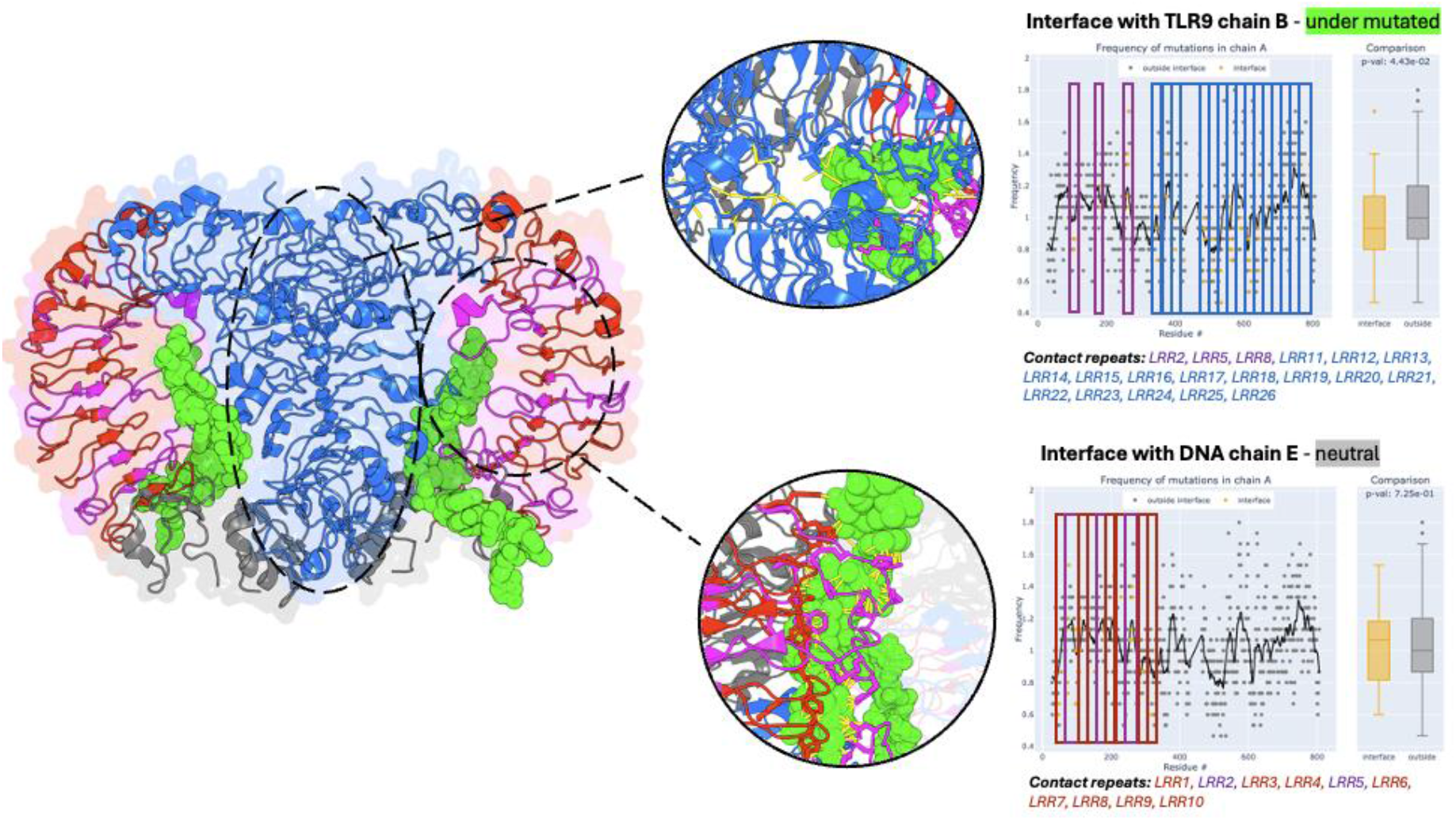
Structural representation of the horse TLR9-DNA complex (PDB: 3WPC (39)) with the distribution of mutations plotted along the length of TLR9. Interactions between LRRs in hTLR5 chains are shown in blue, while those interacting with DNA are shown in red. LRRs involved in both interactions are highlighted in purple. These contact repeats are labeled below the mutation frequency plot and boxed within it. Close-up views of these interactions are circled for clarity. There are two conserved interaction interfaces in TLR9. One interface (bottom close-up), includes the Z-loop binding the single stranded RNA and contributing to the homodimer interface. The second interface binds secondary ligands, including various ssRNA degradation products, but also synthetic agonists and antagonists with therapeutic applications.

## Results

### Sequence divergence of LRR segments

The averages of the features of the individual repeats in all human TLRs are summarized in Table 1, where they are also compared to the features of the archetypal LRR protein, the ribonuclease inhibitor (RNH1). The first sign of the divergence is the varying length of the individual repeats. The standard deviations of repeat lengths in the TLRs are between 4 to 20 times higher than in the RNH1 protein, being the highest in the TLRs with most divergent repeats (TLR7/8/9) due to their long loops between these repeats (Z-loops). The average percent sequence identity between repeats is significantly lower in TLR proteins than in RNH1, indicating greater irregularities among consecutive repeats. Excess length of individual repeats leads to the emergence of “embellishments” - additional loops that sometime form hairpins that break outside of the torus outline. Such elements often bind ligands or participate in the dimerization interfaces, as would be discussed in the next section.

**Table 1.** Overview of repeat divergence in human TLR 1-10 compared against the canonical LRR protein, the ribonuclease inhibitor (PDB:1DFJ). The start and end of repeats were defined by the first conserved leucine position within the “LxL” consensus sequence as discussed in the Methods section. TLRs are colored according to their family. Mean number of conserved positions defined according to the classic consensus sequence pattern: LxLxxN/CxLxxL.

| Identification | Total Number of Repeats | Mean Repeat Length ± SD | Mean Repeat Percent Identity ± SD | Mean Number of Conserved Positions |
| --- | --- | --- | --- | --- |
| <u>Ribonuclease Inhibitor</u> | 14 | 29.5 ± 0.5 | 31.84 ± 11.85 | 5/5 |
| TLR1 | 17 | 25.82 ± 2.81 | 18.5 ± 5.44 | 3.76/5 |
| TLR2 | 19 | 25.79 ± 2.50 | 20.55 ± 5.21 | 4.47/5 |
| TLR3 | 23 | 23.39 ± 2.46 | 24.15 ± 6.00 | 4.61/5 |
| TLR4 | 21 | 25.52 ± 1.99 | 22.53 ± 6.7 | 4.29/5 |
| TLR5 | 21 | 25.90 ± 3.32 | 21.49 ± 6.23 | 4.33/5 |
| TLR6 | 17 | 25.65 ± 2.54 | 19.20 ± 5.62 | 3.94/5 |
| TLR7 | 25 | 29.32 ± 10.09 | 23.43 ± 5.80 | 4.60/5 |
| TLR8 | 25 | 28.96 ± 8.53 | 24.44 ± 6.86 | 4.56/5 |
| TLR9 | 21 | 29.09 ± 8.01 | 24.56 ± 6.97 | 4.48/5 |
| TLR10 | 19 | 25.42 ± 2.39 | 18.34 ± 5.22 | 4.05/5 |

### Local Curvature Analysis

The human genome contains 10 TLR proteins, while other mammals, such as mice, may have up to 13. Figure 2 shows cartoon figures of the structures of the LRR domain of all the 10 human TLRs, 3 of which are AF2 models, with the local curvature of the internal beta sheet and the lengths of the consecutive repeats shown in the graphs below each thumbnail.

As shown on Figure 2, the structures of the LRR domains of all the human TLRs can be described as curved solenoids or toruses. However, even casual observation of the thumbnail figures in Figure 2 shows significant deviations from the ideal torus shape. This is captured by the local curvature measures introduced in the Methods section and shown in the graphs in Figure 2. Interestingly, regions with locally modified curvatures are not located in the same region in different TLRs. In TLR2, 3, and 9, they are located approximately in the middle of the protein, in TLR10 at both ends, and in TLR5 toward the N-terminus. TLR1, 6, 7, and 8 contain long regions with no significant curvature deviations.

Individual LRRs exhibit such a high divergence on the sequence level that some authors suggested that these are non-LRR inserts into the LRR domains (27). This is true for some LRR proteins, especially in bacteria or invertebrates. However, in hTLRs, despite local deviations of the curvature on the concave side of the beta-sheets, the presence of non-canonical residues at some of the usually conserved positions, and sometimes extreme variations in length, the overall LRR signature pattern is conserved and the network of hydrogen bonds in the central beta-sheet is continuous. More careful sequence and phylogenetic analysis, especially between closest divergent paralogs, such as TLR1 and TLR2, or TLR4 and CD180 (data not shown), suggests that the “unusual” repeats arose by divergence rather than non-homologous inserts.

### Functional consequences of the repeat divergence

In the following we show that the divergence of individual repeats in specific TLRs is connected to their unique functions. In the first approximation, we can classify the repeats in each TLR into “structural” repeats with the classical RI class pattern, constant length and local radius vs. “functional” repeats that show significant departure from the classical pattern and varied length and local curvature. Below, we discuss the structural and functional adaptations of individual repeats across TLR subfamilies and demonstrate their relationship to LRR repeat divergence. Additionally, we incorporate data from the human population polymorphism database to demonstrate that the “functional” repeats are still evolving their specific contributions to the TLRs functions.

The vertebrate TLRs are typically divided into 6 subfamilies, differing in their functions and structural features (28). Five of these families are represented in the human genome and an example from each family was analyzed in detail, as described below.

#### TLR1/2/6/10 family

TLR2 binds tri-acylated (in complex with TLR1) or di-acylated (in complex with TLR6) lipopeptides, as well as lipoteichoic acid (LTA), peptidoglycans (PGN), lipoarabinomannan (LAM) and potentially other ligands. Lipopeptides are a broad class of compounds that vary among bacterial species or even between strains, and an individual’s response to specific bacteria may depend on an exact lipopeptide being produced by a given bacterium. The hydrophobic portion of these ligands binds within a cavity in the LRR domain of TLRs—an unusual structural feature setting TLR2 apart from almost all other LRR proteins (29). In contrast, the polar “head” of the lipopeptides is bound to the loops between the LRR repeats. Mutations in these loops may affect its specificity against various bacterial species. Interestingly, despite their high sequence similarity to the TLR2, TLR6 and TLR10 lack any cavity in the LRR domain, and TLR1 contains only a small, partially open cavity. As shown in Figure 3, both the ligand and the TLR1/TLR2 complex interfaces exhibit lower-than-average levels of polymorphism in the human population, suggesting purifying selection pressure on conserving the binding specificity in lipoprotein binding and maintaining the TLR1/TLR2 complex stability.

##### Special structural feature

long loops binding to the head of the lipopeptide and cavities in TLR2, both located in the highly divergent repeats LLR9 to LLR12.

#### TLR3

Endosomal receptor that recognizes viral double-stranded RNA (dsRNA), a key pathogen-associated molecular pattern (PAMP) associated with viral infections. It also recognizes endogenous RNA released from necrotic cells (31) after relocating to the surface of certain immune (dendritic cells and macrophages) and non-immune (Fibroblasts, vascular endothelial cells) cell types. Both ligand interaction interfaces show higher than average levels of polymorphism in the human population, suggesting a potential for diverse immune responses to viral infections among human populations, with different alleles potentially conferring varying susceptibility or severity.

##### Special structural feature

Nucleic acid interaction interface formed by clusters of charged residues engaging dsRNA phosphate backbone. The interaction interfaces are localized on loops between the concave and convex sides of the solenoid in the most divergent repeats.

#### TLR4

Recognizes bacterial lipopolysaccharides (LPS), which is a collective term for components of the outer membrane of gram-negative bacteria (33). LPS from different bacterial species and different strains show large variation, especially in the O antigen “head” (34). LPS is a potent activator of the immune system and excessive amounts of LPS in the blood may trigger a septic shock (35). TLR4, in complex with MD2 protein forms a receptor for the LPS molecules. The conserved Lipid A part of LPS is bound in the cavity in the MD2, while the O antigen head is bound by extended loops in the most divergent repeats in TLR4 (LRR9-13). The O antigen binding loops show higher than average levels of polymorphism, and for instance the best studied TLR4 mutations at positions 299 and 399 are both within the most divergent repeats and at or close to the O antigen binding loops. At the same time, protein-protein interfaces stabilizing the TLR4/MD2 heterotetramer are more conserved, hinting at the importance of maintaining the complex structure for the healthy immune response.

##### Special structural feature

conserved MD2 interaction and dimerization interfaces, loop interacting with LPS O antigen emerging from the cavity in MD2. Variation in the O antigen structure affect binding to the MD2/TLR4 complex and result in dramatic differences in the host immune system responses to different bacteria, such as between *Neisseria meningitidis* (hexa-acylated LPS strongly activates TLR4) and *Helicobacter pylori* (fucosylated LPS reducing TLR4 activation). Polymorphisms in the O antigen binding loops can potentially modulate binding of the different forms of LPS to the TLR4/MD2 complex. However, it must be noted that the experimental results of the TLR4 polymorphism on LPS binding are not consistent and the overall effects of different forms of LPS on human immune response are complicated by the multiple levels and multitude of genes involved in the final response.

The loop in the LRR17 repeat recognizes O antigens, and mutations in this loop can modify TLR4 specificity in responses to different bacteria. This structural feature underscores the adaptability of TLR4 in recognizing and responding to a diverse array of pathogenic challenges, highlighting the evolutionary significance of sequence and structural variability within the LRR domains.

**TLR5 i**s a surface receptor, involved in the detection of bacterial flagellin. The LRR receptor domain of TLR5 binds the D1 “core” domain of flagellin, with two flagellin molecules bound by two TLR5 receptors, thereby stabilizing the dimer in its active form. Like TLR3, the interaction with flagellin mostly involves protruding loops from the “lateral” side, in between the convex and concave side of the LRR solenoid. TLR5 polymorphism is associated with diseases such as altered gut microbiome in IBDs and pneumonia, with both weakened responses leading to increased susceptibility to certain infections and hyper-response leading to chronic inflammation. Like the results for TLR3 and TLR4, the interaction interface with the ligand is highly polymorphic, while the interface between other molecules in the receptor complex is more conserved.

#### TLR 7/8/9

A group of closely homologous endosomal receptors, recognizing viral ssRNA (TLR7), its metabolic products (TLR8) and unmethylated CpG motifs in dsDNA (TLR9). These receptors also recognize various endogenous danger associated molecular patterns (DAMPs) and play a role in many non-infectious diseases. For instance, dysregulation and polymorphism in TLR7 is linked to Systemic Lupus Erythematous (SLE) (38).

As an example from this group, we performed in depth analysis of TLR9. As human TLR9 experimental structure is not available, the horse (39) structure was used for the analysis. With 85% sequence identity and no gaps in the alignments between the human and horse TLR9, we decided to use the horse structure instead of the AF model.

TLR9 recognizes unmethylated CpG DNA motifs, while TLR7 recognizes uridine- and guanosine-rich andTLR8 short uridine-rich sequences. All these motifs are found predominantly in viral and bacterial genomes. However, mitochondrial RNA, self-derived miRNAs, RNA-containing complexes from apoptotic cells and oxidized guanosine nucleotides also contribute to TLR7 activation, particularly in inflammatory and autoimmune conditions.

##### Special structural feature

long loop (Z loop) in the LRR14 position in TLRs 7/8/9 contributes to this subfamily’s large repeat length standard deviation. In LRRs 11-15 form a ligand binding groove for ssRNA binding, with two binding pockets, one focusing on binding ssRNA and second on additional ligands stabilizing the dimerization. Interestingly, in TLR8 the same binding pockets have different binding specificities.

## Discussion

The results of our study highlight the significant structural diversity between the individual repeats in the leucine-rich repeat (LRR) domains of human Toll-like receptor (hTLRs) proteins. This is evident at the sequence level through variations in repeat length and a lack of conservation, even at the positions that typically define the LRR motif. The divergence between sequences of the individual repeats is so large that profile or HHM based sequence analysis programs fail to recognize all the repeats in most LRR proteins and only recent AI based algorithms were able to match the structural analysis approaches. The standard deviation of repeat lengths in TLRs and the level of conservation of the signature sequence pattern is substantially higher than in regular LRR proteins like ribonuclease inhibitor (RNH1) or even paralogs of individual TLRs, such as a TLR4 paralog, CD180. The divergence of the individual repeats extends to the structural level, where despite the conserved pattern of beta strand-loop-alpha helix/irregular structure-loop of each repeat, they differ in the length of the loops, level of irregularity in the convex side of the solenoid and the local curvature on the concave side. This leads to irregular solenoid structures, long loops protruding away from the body of the solenoid and other structural features rarely observed in “classical”, or highly regular LRR proteins. High levels of divergence of individual LRR lead to arguments that in some proteins the repeats are interrupted by non-homologous sections (27). While this is true for some LRR domains, especially in bacteria and invertebrates, the RI class analyzed here have continuous central beta-sheets with conserved hydrogen-bond patterns and recognizable patterns across all repeats, suggesting their common evolutionary history and evolution driven by functional specificity of individual TLRs.

At the same time, structural differences between different TLRs are not captured accurately by the global similarity measures such as RMSD or TM-score. For instance, the RMSD between TLR4 and TLR5, which have different binding mechanisms and functions, have an RMSD of 2.48Å. The functional consequences of variability in repeat length and sequence can be fully appreciated only by analyzing the structural diversity of the repeats. Variations in the local curvature and structural embellishments such as loops and cavities contribute to what we propose to call function-defining regions in the TLRs, which are involved in ligand binding or dimerization interfaces and are essential for the specialized functions of individual TLRs. For example, loops in TLR2 located on a highly divergent LRR12 play a crucial role in binding to the O antigen head of lipoproteins, the natural substrate of TLR2. Similarly, the nucleic acid interaction interface in TLR3 is formed by loops in the highly divergent repeats. In TLR4, loops binding the variable head of the LPS molecules contribute to the specificity of human innate system recognizing different bacterial pathogens. At the same time, polymorphism in the sequences of human TLRs across human population is contributing to the differences in individual responses to pathogens, but also to the prevalence and outcomes of various immune diseases. In our paper we show that patterns of polymorphisms support the role of specific repeats in individual TLRs, generally following the pattern of high variability on the ligand binding regions and conservation in regions involved in maintaining the signaling pathways, such a dimerization interfaces. These findings illustrate the importance of understanding the relationship between sequence divergence, local structural changes, and functional specialization of individual TLRs

Our study provides a comprehensive analysis of the structural and functional diversity within the LRR domains of the human TLRs. The observed variations in repeat length, sequence conservation, and its consequences in local structural features such as curvature and extra loops or cavities highlight the intricate mechanisms by which TLRs have evolved to recognize a wide array of pathogen-associated molecular patterns. This knowledge enhances our understanding of TLR-mediated innate immunity and opens avenues for developing targeted therapeutics that modulate TLR activity.

The LRR domains are not unique to the immune receptors; LRR is a motif found in hundreds of thousands of proteins from all kingdoms of life, typically serving as a ligand binding or protein-protein interaction domain in diverse pathways (6). The human genome codes for about 400 LRR-containing proteins active in different processes, including immune response, apoptosis, autophagy, ubiquitin-related processes, nuclear mRNA transport, and neuronal development (40). Toll-like receptors, a subset of all proteins with LRR domains, are also widely distributed in all kingdoms of life. Direct orthologous lines of the human TLRs can be traced to early vertebrates, but large repertoires of TLRs are also present in most invertebrates, where they emerged by lineage specific expansions. Interestingly, large families of proteins involved in the plant equivalent of innate immunity, the R proteins, have receptor domains homologous to the LRR domains of TLRs. As the binding specificities are likely to at least partly overlap between TLRs of different species that deal with similar pathogens, it is possible that similar function driven solutions have emerged by parallel evolution, which makes analyses like those presented here applicable to a much larger sets of proteins.

## Supporting information

overview of the supplemental materials

Dataset D2 and Figures S1 and S2: comparison of the experimental and AF models

Dataset D1: Local curvature and lengths of individual repeats

Dataset D3: exact positions of individual repeats in human TLRs

## Acronyms

TLRs: Toll-like receptors
LRRs: Leucine Rich Repeats
TIR domain: Toll/interleukin-1 receptor domain
PAMPs: Pathogen-Associated Molecular Patterns
DAMPs: Damage (or Danger) -Associated Molecular Patterns

## References

1. N. W. Palm, R. Medzhitov, Pattern recognition receptors and control of adaptive immunity. Immunol Rev 227, 221–233 (2009).

2. T. Duan, Y. Du, C. Xing, H. Y. Wang, R. F. Wang, Toll-Like Receptor Signaling and Its Role in Cell-Mediated Immunity. Front Immunol 13, 812774 (2022).

3. M. A. Delgado, R. A. Elmaoued, A. S. Davis, G. Kyei, V. Deretic, Toll-like receptors control autophagy. EMBO J 27, 1110–1121 (2008).

4. Y. Yang et al., Toll-like receptors: Triggers of regulated cell death and promising targets for cancer therapy. Immunol Lett 223, 1–9 (2020).

5. M. A. Sanjuan et al., Toll-like receptor signalling in macrophages links the autophagy pathway to phagocytosis. Nature 450, 1253–1257 (2007).

6. B. Kobe, A. V. Kajava, The leucine-rich repeat as a protein recognition motif. Curr Opin Struct Biol 11, 725–732 (2001).

7. E. I. Deryusheva, A. V. Machulin, O. V. Galzitskaya, Diversity and features of proteins with structural repeats. Biophysical Reviews 15, 1159–1169 (2023).

8. D. Kaur, S. Patiyal, N. Sharma, S. S. Usmani, G. P. S. Raghava, PRRDB 2.0: a comprehensive database of pattern-recognition receptors and their ligands. Database (Oxford) 2019 (2019).

9. Y. Belkhadir, R. Subramaniam, J. L. Dangl, Plant disease resistance protein signaling: NBS-LRR proteins and their partners. Curr Opin Plant Biol 7, 391–399 (2004).

10. L. Yu, L. Wang, S. Chen, Endogenous toll-like receptor ligands and their biological significance. Journal of cellular and molecular medicine 14, 2592–2603 (2010).

11. Z. Chang, Important aspects of Toll-like receptors, ligands and their signaling pathways. Inflammation research 59, 791–808 (2010).

12. O.K. Tørresen et al., Tandem repeats lead to sequence assembly errors and impose multi-level challenges for genome and protein databases. Nucleic acids research 47, 10994–11006 (2019).

13. M. G. Tassia, N. V. Whelan, K. M. Halanych, Toll-like receptor pathway evolution in deuterostomes. Proceedings of the National Academy of Sciences 114, 7055–7060 (2017).

14. B. Kobe, J. Deisenhofer, A structural basis of the interactions between leucine-rich repeats and protein ligands. Nature 374, 183–186 (1995).

15. M. Pellegrini, Tandem Repeats in Proteins: Prediction Algorithms and Biological Role. Front Bioeng Biotechnol 3, 143 (2015).

16. T. Hrabe, A. Godzik, ConSole: using modularity of contact maps to locate solenoid domains in protein structures. BMC Bioinformatics 15, 119 (2014).

17. InterPro (2025) Ribonuclease inhibitor (RINI_HUMAN, P13489).

18. K. Qiu, S. Dunin-Horkawicz, A. Lupas, Exploiting protein language model sequence representations for repeat detection. bioRxiv 10.1101/2024.06.07.596093, 2024.2006.2007.596093 (2024).

19. K. Ichii, A. Takkouche, e. al., Protein Repeat Recognition Through Unsupervised Clustering of Protein Large Language Model Embeddings. Bioinformatics in preparation (2024).

20. Y. Zhang, J. Skolnick, Scoring function for automated assessment of protein structure template quality. Proteins 57, 702–710 (2004).

21. S. Das, N. L. Dawson, C. A. Orengo, Diversity in protein domain superfamilies. Curr Opin Genet Dev 35, 40–49 (2015).

22. Y. Nedelec et al., Genetic Ancestry and Natural Selection Drive Population Differences in Immune Responses to Pathogens. Cell 167, 657–669 e621 (2016).

23. P. Strzelecki, H. Von Der Mosel, Integral Menger curvature for surfaces. Advances in Mathematics 226, 2233–2304 (2011).

24. S. K. Burley et al., RCSB Protein Data Bank (RCSB.org): delivery of experimentally-determined PDB structures alongside one million computed structure models of proteins from artificial intelligence/machine learning. Nucleic Acids Res 51, D488–D508 (2023).

25. B. Kobe, J. Deisenhofer, Mechanism of ribonuclease inhibition by ribonuclease inhibitor protein based on the crystal structure of its complex with ribonuclease A. Journal of molecular biology 264, 1028–1043 (1996).

26. S. Gudmundsson et al., Variant interpretation using population databases: Lessons from gnomAD. Human mutation 43, 1012–1030 (2022).

27. N. Matsushima et al., Analyses of non-leucine-rich repeat (non-LRR) regions intervening between LRRs in proteins. Biochim Biophys Acta 1790, 1217–1237 (2009).

28. J. C. Roach et al., The evolution of vertebrate Toll-like receptors. Proceedings of the National Academy of Sciences 102, 9577–9582 (2005).

29. R. Namou, L. Jaroszewski, K. Ichii, A. Takkouche, A. Godzik, Homology vs. homoplasy in innate receptor ligand recognition: the story of the TLR2 cavity. in preparation (2025).

30. M. S. Jin et al., Crystal structure of the TLR1-TLR2 heterodimer induced by binding of a tri-acylated lipopeptide. Cell 130, 1071–1082 (2007).

31. K. Kariko, H. Ni, J. Capodici, M. Lamphier, D. Weissman, mRNA is an endogenous ligand for Toll-like receptor 3. J Biol Chem 279, 12542–12550 (2004).

32. C. S. Lim et al., TLR3 forms a highly organized cluster when bound to a poly(I:C) RNA ligand. Nat Commun 13, 6876 (2022).

33. M. Caroff, D. Karibian, Structure of bacterial lipopolysaccharides. Carbohydrate research 338, 2431–2447 (2003).

34. I. Lerouge, J. Vanderleyden, O-antigen structural variation: mechanisms and possible roles in animal/plant–microbe interactions. FEMS microbiology reviews 26, 17–47 (2002).

35. J. A. Kellum, C. Ronco, The role of endotoxin in septic shock. Crit Care 27, 400 (2023).

36. K. Park et al., Control of repeat-protein curvature by computational protein design. Nature structural & molecular biology 22, 167–174 (2015).

37. S. I. Yoon et al., Structural basis of TLR5-flagellin recognition and signaling. Science 335, 859–864 (2012).

38. Y. H. Lee, S. Choi, J. Ji, G. Song, Association between toll-like receptor polymorphisms and systemic lupus erythematosus: a meta-analysis update. Lupus 25, 593–601 (2016).

39. U. Ohto et al., Structural basis of CpG and inhibitory DNA recognition by Toll-like receptor 9. Nature 520, 702–705 (2015).

40. A. C. Ng et al., Human leucine-rich repeat proteins: a genome-wide bioinformatic categorization and functional analysis in innate immunity. Proceedings of the National Academy of Sciences 108, 4631–4638 (2011).

