## Supplementary material for "Divergence of the individual repeats in the leucine-rich repeat domains of human Toll-like receptors explain their diversity and functional adaptations": overview of the supplemental materials

**This PDF file includes:**

Description of datasets DS1 to DS3

**Other supporting materials for this manuscript include the following:**

Datasets DS1 to DS3

**Supporting Information** contains three datasets of tables and figures:

**Dataset S1:** local measurements of hTLR1-10, ribonuclease inhibitor, and CD180 structures. Curvature (circumradius), repeat lengths calculated from experimental structures where available or AlphaFold models where they were not.

**Dataset S2**. Comparison of local curvature and circumcenter values for individual repeats in human ribonuclease Inhibitor (PDB:2BNH) and receptor domain of hTLR1 as calculated for the experimental structures, (PDB IDs 2BNH and 6NIH, respectively) and their AlphaFold2 and ESM2 models. Each table is followed by a Figure, S1 and S2, respectively, providing visual comparison of the circumcenter values for the three models.

**Dataset S3:** positions (start and end) for individual LRR repeats for hTLR1–10, ribonuclease inhibitor, and CD180, with repeat lengths shown in parentheses.
