## Supplementary material for "Divergence of the individual repeats in the leucine-rich repeat domains of human Toll-like receptors explain their diversity and functional adaptations": Dataset D2 and Figures S1 and S2: comparison of the experimental and AF models

**Ribonuclease Inhibitor (PDB:2BNH): *Experimental***

| **Res. 1** | **Res. 2** | **Res. 3** | **Curvature** | **Circumradius** | **Repeat Length 1-2** | **Repeat Length 2-3** |
| --- | --- | --- | --- | --- | --- | --- |
| V28 | L56 | L85 | 0.0706 | 14.17 | 28 | 29 |
| L56 | L85 | L113 | 0.0428 | 23.39 | 29 | 28 |
| L85 | L113 | L142 | 0.053 | 18.88 | 28 | 29 |
| L113 | L142 | L170 | 0.0425 | 23.52 | 29 | 28 |
| L142 | L170 | L199 | 0.0858 | 11.65 | 28 | 29 |
| L170 | L199 | L227 | 0.059 | 16.94 | 29 | 28 |
| L199 | L227 | L256 | 0.0375 | 26.67 | 28 | 29 |
| L227 | L256 | L284 | 0.0674 | 14.84 | 29 | 28 |
| L256 | L284 | L313 | 0.0426 | 23.45 | 28 | 29 |
| L284 | L313 | L341 | 0.0639 | 15.64 | 29 | 28 |
| L313 | L341 | L370 | 0.0517 | 19.33 | 28 | 29 |
| L341 | L370 | L398 | 0.0742 | 13.48 | 29 | 28 |
| L370 | L398 | L427 | 0.0356 | 28.07 | 28 | 29 |

**Ribonuclease Inhibitor (PDB:2BNH): *AlphaFold2***

| **Res. 1** | **Res. 2** | **Res. 3** | **Curvature** | **Circumradius** | **Repeat Length 1-2** | **Repeat Length 2-3** |
| --- | --- | --- | --- | --- | --- | --- |
| V28 | L56 | L85 | 0.0713 | 14.03 | 28 | 29 |
| L56 | L85 | L113 | 0.0335 | 29.88 | 29 | 28 |
| L85 | L113 | L142 | 0.0632 | 15.82 | 28 | 29 |
| L113 | L142 | L170 | 0.0661 | 15.12 | 29 | 28 |
| L142 | L170 | L199 | 0.0884 | 11.31 | 28 | 29 |
| L170 | L199 | L227 | 0.0534 | 18.73 | 29 | 28 |
| L199 | L227 | L256 | 0.0489 | 20.44 | 28 | 29 |
| L227 | L256 | L284 | 0.05 | 20 | 29 | 28 |
| L256 | L284 | L313 | 0.0545 | 18.34 | 28 | 29 |
| L284 | L313 | L341 | 0.0666 | 15.01 | 29 | 28 |
| L313 | L341 | L370 | 0.0672 | 14.88 | 28 | 29 |
| L341 | L370 | L398 | 0.0587 | 17.04 | 29 | 28 |
| L370 | L398 | L427 | 0.0638 | 15.68 | 28 | 29 |

**Ribonuclease Inhibitor (PDB:2BNH): *ESM***

| **Res. 1** | **Res. 2** | **Res. 3** | **Curvature** | **Circumradius** | **Repeat Length 1-2** | **Repeat Length 2-3** |
| --- | --- | --- | --- | --- | --- | --- |
| V28 | L56 | L85 | 0.0665 | 15.04 | 28 | 29 |
| L56 | L85 | L113 | 0.0319 | 31.39 | 29 | 28 |
| L85 | L113 | L142 | 0.0576 | 17.35 | 28 | 29 |
| L113 | L142 | L170 | 0.0546 | 18.32 | 29 | 28 |
| L142 | L170 | L199 | 0.0785 | 12.74 | 28 | 29 |
| L170 | L199 | L227 | 0.0552 | 18.12 | 29 | 28 |
| L199 | L227 | L256 | 0.0537 | 18.63 | 28 | 29 |
| L227 | L256 | L284 | 0.0486 | 20.58 | 29 | 28 |
| L256 | L284 | L313 | 0.0624 | 16.03 | 28 | 29 |
| L284 | L313 | L341 | 0.0603 | 16.58 | 29 | 28 |
| L313 | L341 | L370 | 0.0706 | 14.16 | 28 | 29 |
| L341 | L370 | L398 | 0.0571 | 17.52 | 29 | 28 |
| L370 | L398 | L427 | 0.0623 | 16.06 | 28 | 29 |

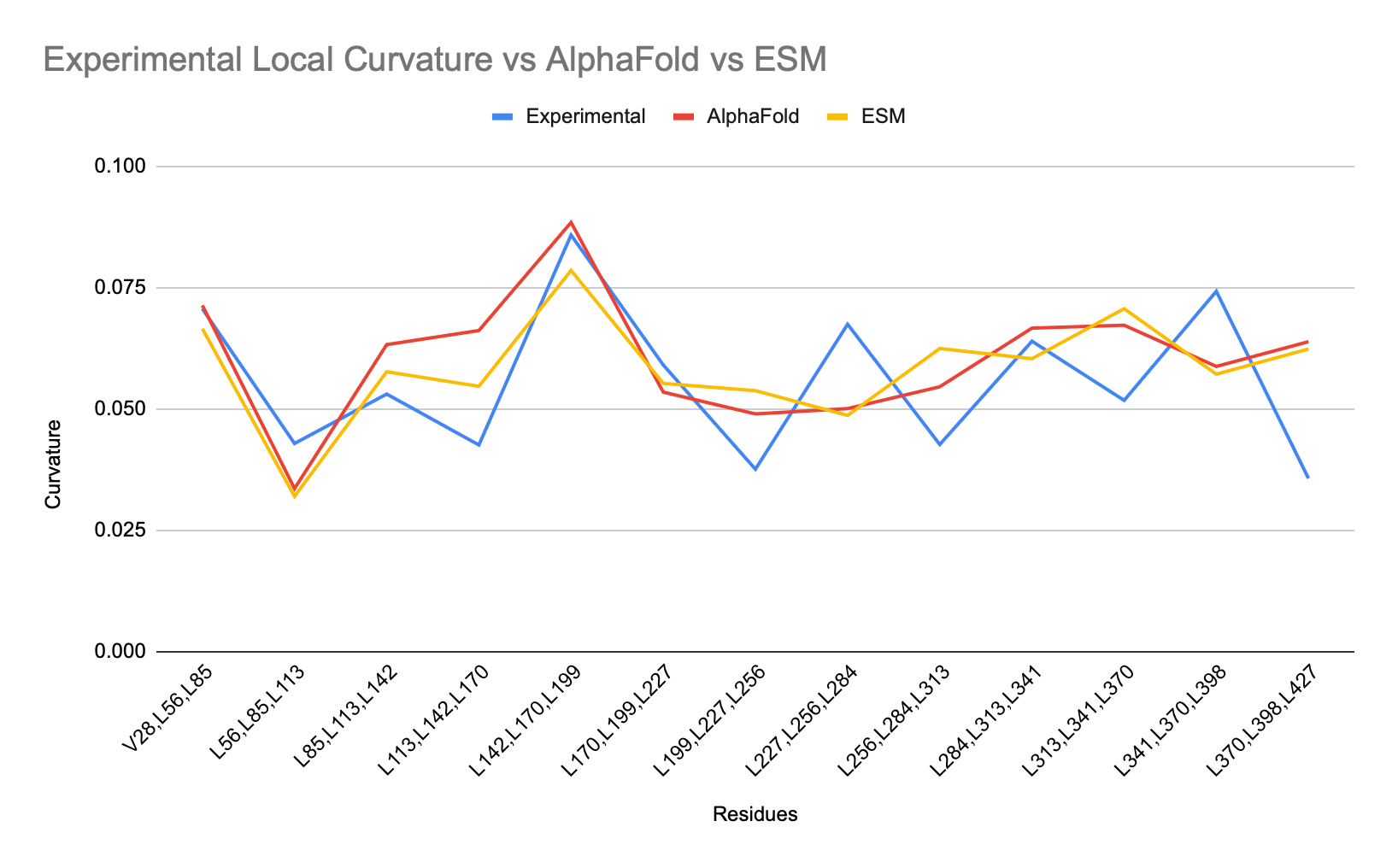

***Human TLR1* (PDB:6NIH): *Experimental***

| **Res. 1** | **Res. 2** | **Res. 3** | **Curvature** | **Circumradius** | **Repeat Length 1-2** | **Repeat Length 2-3** |
| --- | --- | --- | --- | --- | --- | --- |
| L50 | L74 | L98 | 0.0387 | 25.86 | 24 | 24 |
| L74 | L98 | L119 | 0.031 | 32.21 | 24 | 21 |
| L98 | L119 | L144 | 0.062 | 16.14 | 21 | 25 |
| L119 | L144 | V167 | 0.0749 | 13.36 | 25 | 23 |
| L144 | V167 | L192 | 0.01 | 99.54 | 23 | 25 |
| V167 | L192 | L216 | 0.0471 | 21.22 | 25 | 24 |
| L192 | L216 | L249 | 0.0655 | 15.26 | 24 | 33 |
| L216 | L249 | F276 | 0.0427 | 23.41 | 33 | 27 |
| L249 | F276 | L302 | 0.0278 | 35.97 | 27 | 26 |
| F276 | L302 | F331 | 0.0308 | 32.42 | 26 | 29 |
| L302 | F331 | L353 | 0.0606 | 16.49 | 29 | 22 |
| F331 | L353 | L377 | 0.0166 | 60.13 | 22 | 24 |
| L353 | L377 | L403 | 0.0453 | 22.09 | 24 | 26 |
| L377 | L403 | L428 | 0.0441 | 22.69 | 26 | 25 |
| L403 | L428 | L450 | 0.0534 | 18.74 | 25 | 22 |
| L428 | L450 | L473 | 0.0373 | 26.84 | 22 | 23 |

***Human TLR1* (PDB:6NIH): *AlphaFold2***

| ***Res. 1*** | ***Res. 2*** | ***Res. 3*** | ***Curvature*** | ***Circumradius*** | ***Repeat Length 1-2*** | ***Repeat Length 2-3*** |
| --- | --- | --- | --- | --- | --- | --- |
| *L50* | *L74* | *L98* | *0.0373* | *26.82* | *24* | *24* |
| *L74* | *L98* | *L119* | *0.0366* | *27.33* | *24* | *21* |
| *L98* | *L119* | *L144* | *0.0684* | *14.63* | *21* | *25* |
| *L119* | *L144* | *V167* | *0.0588* | *17* | *25* | *23* |
| *L144* | *V167* | *L192* | *0.0154* | *64.75* | *23* | *25* |
| *V167* | *L192* | *L216* | *0.053* | *18.88* | *25* | *24* |
| *L192* | *L216* | *L249* | *0.0509* | *19.65* | *24* | *33* |
| *L216* | *L249* | *F276* | *0.0408* | *24.53* | *33* | *27* |
| *L249* | *F276* | *L302* | *0.0167* | *59.77* | *27* | *26* |
| *F276* | *L302* | *F331* | *0.0472* | *21.18* | *26* | *29* |
| *L302* | *F331* | *L353* | *0.0484* | *20.66* | *29* | *22* |
| *F331* | *L353* | *L377* | *0.0383* | *26.08* | *22* | *24* |
| *L353* | *L377* | *L403* | *0.0458* | *21.82* | *24* | *26* |
| *L377* | *L403* | *L428* | *0.0327* | *30.54* | *26* | *25* |
| *L403* | *L428* | *L450* | *0.0406* | *24.61* | *25* | *22* |
| *L428* | *L450* | *L473* | *0.0478* | *20.9* | *22* | *23* |

***Human TLR1* (PDB:6NIH): *ESM***

| **Res. 1** | **Res. 2** | **Res. 3** | **Curvature** | **Circumradius** | **Repeat Length 1-2** | **Repeat Length 2-3** |
| --- | --- | --- | --- | --- | --- | --- |
| L50 | L74 | L98 | 0.0361 | 27.73 | 24 | 24 |
| L74 | L98 | L119 | 0.0333 | 30.05 | 24 | 21 |
| L98 | L119 | L144 | 0.0647 | 15.45 | 21 | 25 |
| L119 | L144 | V167 | 0.0665 | 15.03 | 25 | 23 |
| L144 | V167 | L192 | 0.0122 | 82.15 | 23 | 25 |
| V167 | L192 | L216 | 0.0544 | 18.37 | 25 | 24 |
| L192 | L216 | L249 | 0.0447 | 22.38 | 24 | 33 |
| L216 | L249 | F276 | 0.0435 | 22.97 | 33 | 27 |
| L249 | F276 | L302 | 0.0206 | 48.52 | 27 | 26 |
| F276 | L302 | F331 | 0.0518 | 19.3 | 26 | 29 |
| L302 | F331 | L353 | 0.0488 | 20.49 | 29 | 22 |
| F331 | L353 | L377 | 0.0359 | 27.83 | 22 | 24 |
| L353 | L377 | L403 | 0.0425 | 23.54 | 24 | 26 |
| L377 | L403 | L428 | 0.033 | 30.26 | 26 | 25 |
| L403 | L428 | L450 | 0.0514 | 19.44 | 25 | 22 |
| L428 | L450 | L473 | 0.0356 | 28.1 | 22 | 23 |

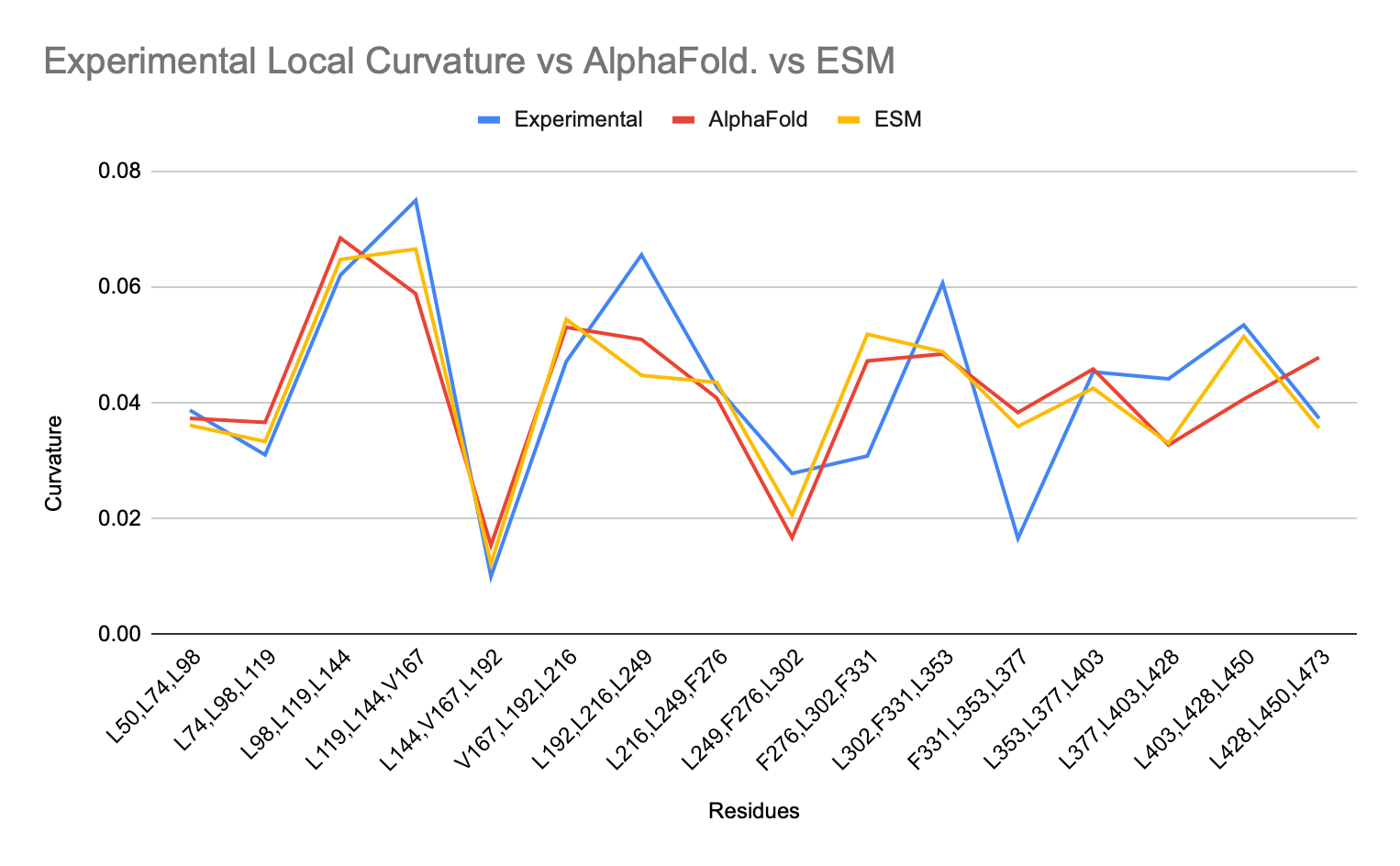
