## Supplementary material for "Divergence of the individual repeats in the leucine-rich repeat domains of human Toll-like receptors explain their diversity and functional adaptations": Dataset D3: exact positions of individual repeats in human TLRs

**Classic Consensus Sequence: LxLxxN/CxLxxL**

**TLR1** *(manually measured from ladder positions)*

>LRR1  50-74       **L**N**I**SQ**N**Y**I**SE**L**WTSDILSLSKLRIL (25)

>LRR2  74-98       **L**I**I**SH**N**R**I**QY**L**DISVFKFNQELEYL (25)

>LRR3  98-119      **L**D**L**SH**N**K**L**VK**I**SCHPTVNLKHL (22)

>LRR4  119-144     **L**D**L**SF**N**A**F**DA**L**PICKEFGNMSQLKFL (26)

>LRR5  144-167     **L**G**L**STTH**L**EKSSVLPIAHLNISKV (24)

>LRR6  167-192     **V**L**L**VLGETYGEKEDPEGLQDFNTESL (26)

>LRR7  192-216     **L**H**I**VFPTNKE**F**HFILDVSVKTVANL (25)

>LRR8  216-249     **L**E**L**SNIKCVLEDNKCSYFLSILAKLQTNPKLSNL (34)

>LRR9  249-276     **L**T**L**NNIETTWNSFIRILQLVWHTTVWYF (28)

>LRR10 276-302     **F**S**I**SNVK**L**QGQLDFRDFDYSGTSLKAL (27)

>LRR11 302-331     **L**S**I**HQVVSDV**F**GFPQSYIYEIFSNMNIKNF (30)

>LRR12 331-353     **F**T**V**SGTR**M**VH**M**LCPSKISPFLHL (23)

>LRR13 353-377     **L**D**F**SN**N**L**L**TDTVFENCGHLTELETL (24)

>LRR14 377-403     **L**I**L**QM**N**Q**L**KE**L**SKIAEMTTQMKSLQQL (27)

>LRR15 403-428     **L**D**I**SQ**N**S**V**SYDEKKGDCSWTKSLLSL (26)

>LRR16 428-450     **L**N**M**SS**N**I**L**TDTIFRCLPPRIKVL (23)

>LRR17 450-473     **L**D**L**HS**N**K**I**KS**I**PKQVVKLEALQEL (24)

**TLR2** *(manually measured from ladder positions)*

>LRR1 57-81        **L**D**L**SN**N**R**I**TY**I**SNSDLQRCVNLQAL (25)

>LRR2 81-105       **L**V**L**TS**N**G**I**NT**I**EEDSFSSLGSLEHL (25)

>LRR3 105-129      **L**D**L**SY**N**Y**L**SN**L**SSSWFKPLSSLTFL (25)

>LRR4 129-154      **L**N**L**LG**N**P**Y**KT**L**GETSLFSHLTKLQIL (26)

>LRR5 154-179      **L**R**V**GNMDTFTKIQRKDFAGLTFLEEL (26)

>LRR6 179-203      **L**E**I**DA**S**D**L**QS**Y**EPKSLKSIQNVSHL (25)

>LRR7 203-227      **L**I**L**HMKQHIL**L**LEIFVDVTSSVECL (25)

>LRR8 227-254      **L**E**L**RD**T**D**L**DT**F**HFSELSTGETNSLIKKF (28)

>LRR9 254-282      **F**T**F**RNVK**I**TDESLFQVMKLLNQISGLLEL (29)

>LRR10 282-312     **L**E**F**DD**C**T**L**NG**V**GNFRASDNDRVIDPGKVETL (31)

>LRR11 312-341     **L**T**I**RRLH**I**PR**F**YLFYDLSTLYSLTERVKRI (30)

>LRR12 341-365     **I**T**V**ENSK**V**FL**V**PCLLSQHLKSLEYL (25)

>LRR13 365-392     **L**D**L**SE**N**L**M**VEEYLKNSACEDAWPSLQTL (28)

>LRR14 392-418     **L**I**L**RQ**N**H**L**AS**L**EKTGETLLTLKNLTNI (27)

>LRR15 418-441     **I**D**I**SK**N**S**F**HS**M**PETCQWPEKMKYL (24)

>LRR16 441-462     **L**N**L**SS**T**R**I**HS**V**TGCIPKTLEIL (22)

>LRR17 462-482     **L**D**V**SN**N**N**L**NL**F**SLNLPQLKEL (21)

>LRR18 482-504     **L**Y**I**SR**N**K**L**MT**L**PDASLLPMLLVL (23)

>LRR19 504-528     **L**K**I**SR**N**Q**L**KS**V**PDGIFDRLTSLQKI (25)

**TLR3** *(manually measured from ladder positions)*

>LRR1 56-80        **L**N**L**TH**N**Q**L**RR**L**PAANFTRYSQLTSL (25)

>LRR2 80-104       **L**D**V**GF**N**T**I**SK**L**EPELCQKLPMLKVL (25)

>LRR3 104-128      **L**N**L**QH**N**E**L**SQ**L**SDKTFAFCTNLTEL (25)

>LRR4 128-152      **L**H**L**MS**N**S**I**QK**I**KNNPFVKQKNLITL (25)

>LRR5 152-176      **L**D**L**SH**N**G**L**SSTKLGTQVQLENLQEL (25)

>LRR6 176-202      **L**L**L**SN**N**K**I**QA**L**KSEELDIFANSSLKKL (27)

>LRR7 202-226      **L**E**L**SS**N**Q**I**KE**F**SPGCFHAIGRLFGL (25)

>LRR8 226-253      **L**F**L**NNVQ**L**GPSLTEKLCLELANTSIRNL (28)

>LRR9 253-279      **L**S**L**SN**S**Q**L**ST**T**SNTTFLGLKWTNLTML (27)

>LRR10 279-303     **L**D**L**SY**N**N**L**NV**V**GNDSFAWLPQLEYF (25)

>LRR11 303-327     **F**F**L**EY**N**N**I**QH**L**FSHSLHGLFNVRYL (25)

>LRR12 327-360     **L**N**L**KR**S**FTKQSISLASLPKIDDFSFQWLKCLEHL (34)

>LRR13 360-384     **L**N**M**ED**N**D**I**PG**I**KSNMFTGLINLKYL (25)

>LRR14 384-412     **L**S**L**SN**S**FTSLRTLTNETFVSLAHSPLHIL (29)

>LRR15 412-436     **L**N**L**TK**N**K**I**SK**I**ESDAFSWLGHLEVL (25)

>LRR16 436-461     **L**D**L**GL**N**E**I**GQELTGQEWRGLENIFEI (26)

>LRR17 461-485     **I**Y**L**SY**N**K**Y**LQ**L**TRNSFALVPSLQRL (25)

>LRR18 485-511     **L**M**L**RRVA**L**KN**V**DSSPSPFQPLRNLTIL (27)

>LRR19 511-535     **L**D**L**SN**N**N**I**AN**I**NDDMLEGLEKLEIL (25)

>LRR20 535-567     **L**D**L**QH**N**N**L**AR**L**WKHANPGGPIYFLKGLSHLHIL (33)

>LRR21 567-591     **L**N**L**ES**N**G**F**DE**I**PVEVFKDLFELKII (25)

>LRR22 591-615     **I**D**L**GL**N**N**L**NT**L**PASVFNNQVSLKSL (25)

>LRR23 615-640     **L**N**L**QK**N**L**I**TS**V**EKKVFGPAFRNLTEL (26)

**TLR4** *(manually measured from ladder positions)*

```
>LRR1 59-83        LDLSFNPLRHLGSYSFFSFPELQVL (25)
>LRR2 83-107       LDLSRCEIQTIEDGAYQSLSHLSTL (25)
>LRR3 107-131      LILTGNPIQSLALGAFSGLSSLQKL (25)
>LRR4 131-155      LVAVETNLASLENFPIGHLKTLKEL (25)
>LRR5 155-180      LNVAHNLIQSFKLPEYFSNLTNLEHL (26)
>LRR6 180-208      LDLSSNKIQSIYCTDLRVLHQMPLLNLSL (29)
>LRR7 208-231      LDLSLNPMNFIQPGAFKEIRLHKL (24)
>LRR8 231-258      LTLRNNFDSLNVMKTCIQGLAGLEVHRL (28)
>LRR9 258-288      LVLGEFRNEGNLEKFDKSALEGLCNLTIEEF (31)
>LRR10 288-313     FRLAYLDYYLDDIIDLFNCLTNVSSF (26)
>LRR11 313-335     FSLVSVTIERVKDFSYNFGWQHL (23)
>LRR12 335-356     LELVNCKFGQFPTLKLKSLKRL (22)
>LRR13 356-378     LTFTSNKGGNAFSEVDLPSLEFL (23)
>LRR14 378-404     LDLSRNGLSFKGCCSQSDFGTTSLKYL (27)
>LRR15 404-427     LDLSFNGVITMSSNFLGLEQLEHL (24)
>LRR16 427-452     LDFQHSNLKQMSEFSVFLSLRNLIYL (26)
>LRR17 452-476     LDISHTHTRVAFNGIFNGLSSLEVL (25)
>LRR18 476-501     LKMAGNSFQENFLPDIFTELRNLTFL (26)
>LRR19 501-525     LDLSQCQLEQLSPTAFNSLSSLQVL (25)
>LRR20 525-549     LNMSHNNFFSLDTFPYKCLNSLQVL (25)
>LRR21 549-574     LDYSLNHIMTSKKQELQHFPSSLAFL (26)
TLR5 (manually measured from ladder positions)
>LRR1 51-75        LLLSFNYIRTVTASSFPFLEQLQLL (25)
>LRR2 75-100       LELGSQYTPLTIDKEAFRNLPNLRIL (26)
>LRR3 100-124      LDLGSSKIYFLHPDAFQGLFHLFEL (25)
>LRR4 124-150      LRLYFCGLSDAVLKDGYFRNLKALTRL (27)
>LRR5 150-175      LDLSKNQIRSLYLHPSFGKLNSLKSI (26)
>LRR6 175-203      IDFSSNQIFLVCEHELEPLQGKTLSFFSL (29)
>LRR7 203-231      LAANSLYSRVSVDWGKCMNPFRNMVLEIL (29)
>LRR8 231-258      LDVSGNGWTVDITGNFSNAISKSQAFSL (28)
>LRR9 258-293      LILAHHIMGAGFGFHNIKDPDQNTFAGLARSSVRHL (36)
>LRR10 293-317     LDLSHGFVFSLNSRVFETLKDLKVL (25)
>LRR11 317-341     LNLAYNKINKIADEAFYGLDNLQVL (25)
>LRR12 341-365     LNLSYNLLGELYSSNFYGLPKVAYI (25)
>LRR13 365-389     IDLQKNHIAIIQDQTFKFLEKLQTL (25)
>LRR14 389-408     LDLRDNALTTIHFIPSIPDI (20)
>LRR15 408-428     IFLSGNKLVTLPKINLTANLI (21)
>LRR16 428-453     IHLSENRLENLDILYFLLRVPHLQIL (26)
>LRR17 453-478     LILNQNRFSSCSGDQTPSENPSLEQL (26)
>LRR18 478-502     LFLGENMLQLAWETELCWDVFEGLS (25)
>LRR19 502-531     SHLQVLYLNHNYLNSLPPGVFSHLTALRGL (30)
>LRR20 531-553     LSLNSNRLTVLSHNDLPANLEIL (23)
>LRR21 553-574     LDISRNQLLAPNPDVFVSLSVL (22)

TLR6 (manually measured from ladder positions)
>LRR1 81-105       LRLSHNRIQLLDLSVFKFNQDLEYL (25)
>LRR2 105-126      LDLSHNQLQKISCHPIVSFRHL (22)
>LRR3 126-151      LDLSFNDFKALPICKEFGNLSQLNFL (26)
>LRR4 151-174      LGLSAMKLQKLDLLPIAHLHLSYI (24)
>LRR5 174-199      ILLDLRNYYIKENETESLQILNAKTL (26)
>LRR6 199-223      LHLVFHPTSLFAIQVNISVNTLGCL (25)
>LRR7 223-254      LQLTNIKLNDDNCQVFIKFLSELTRGSTLLNF (32)
>LRR8 254-281      FTLNHIETTWKCLVRVFQFLWPKPVEYL (28)
>LRR9 281-307      LNIYNLTIIESIREEDFTYSKTTLKAL (27)
>LRR10 307-336     LTIEHITNQVFLFSQTALYTVFSEMNIMML (30)
>LRR11 336-358     LTISDTPFIHMLCPHAPSTFKFL (23)
>LRR12 358-382     LNFTQNVFTDSIFEKCSTLVKLETL (25)
>LRR13 382-408     LILQKNGLKDLFKVGLMTKDMPSLEIL (27)
>LRR14 408-433     LDVSWNSLESGRHKENCTWVESIVVL (26)
>LRR15 433-455     LNLSSNMLTDSVFRCLPPRIKVL (23)
>LRR16 455-478     LDLHSNKIKSVPKQVVKLEALQEL (24)
>LRR17 478-500     LNVAFNSLTDLPGCGSFSSLSVL (23)


TLR7 (manually measured from ladder positions)
Reference:                 LxLxxNxLxxL
>LRR1 70-94       LTLTINHIPDISPASFHRLDHLVEI (25)
>LRR2 94-132      IDFRCNCVPIPLGSKNNMCIKRLQIKPRSFSGLTYLKSL (39)
>LRR3 132-153     LYLDGNQLLEIPQGLPPSLQLL (22)
>LRR4 153-177     LSLEANNIFSIRKENLTELANIEIL (25)
>LRR5 177-209     LYLGQNCYYRNPCYVSYSIEKDAFLNLTKLKVL (33)
>LRR6 209-230     LSLKDNNVTAVPTVLPSTLTEL (22)
>LRR7 230-254     LYLYNNMIAKIQEDDFNNLNQLQIL (25)
>LRR8 254-295     LDLSGNCPRCYNAPFPCAPCKNNSPLQIPVNAFDALTELKVL (42)
>LRR9 295-319     LRLHSNSLQHVPPRWFKNINKLQEL (25)
>LRR10 319-345    LDLSQNFLAKEIGDAKFLHFLPSLIQL (27)
>LRR11 345-375    LDLSFNFELQVYRASMNLSQAFSSLKSLKIL (31)
>LRR12 375-402    LRIRGYVFKELKSFNLSPLHNLQNLEVL (28)
>LRR13 402-426    LDLGTNFIKIANLSMFKQFKRLKVI (25)
>LRR14 426-498    IDLSVNKISPSGDSSEVGFCSNARTSVESYEPQVLEQLHYFRYDKY ARSCRFKNKEASFMSVNESCYKYGQTL (73)
>LRR15 498-522    LDLSKNSIFFVKSSDFQHLSFLKCL (25)
>LRR16 522-547    LNLSGNLISQTLNGSEFQPLAELRYL (26)
>LRR17 547-571    LDFSNNRLDLLHSTAFEELHKLEVL (25)
>LRR18 571-601    LDISSNSHYFQSEGITHMLNFTKNLKVLQKL (31)
>LRR19 601-624    LMMNDNDISSSTSRTMESESLRTL (24)
>LRR20 624-655    LEFRGNHLDVLWREGDNRYLQLFKNLLKLEEL (32)
>LRR21 655-680    LDISKNSLSFLPSGVFDGMPPNLKNL (26)
>LRR22 680-704    LSLAKNGLKSFSWKKLQCLKNLETL (25)
>LRR23 704-728    LDLSHNQLTTVPERLSNCSRSLKNL (25)
>LRR24 728-752    LILKNNQIRSLTKYFLQDAFQLRYL (25)
>LRR25 752-778    LDLSSNKIQMIQKTSFPENVLNNLKML


TLR8 (manually measured from ladder positions)
>LRR1 68-92        LDLSDNFITHITNESFQGLQNLTKI (25)
>LRR2 92-130       INLNHNPNVQHQNGNPGIQSNGLNITDGAFLNLKNLREL (39)
>LRR3 130-151      LLLEDNQLPQIPSGLPESLTEL (22)
>LRR4 151-175      LSLIQNNIYNITKEGISRLINLKNL (25)
>LRR5 175-206      LYLAWNCYFNKVCEKTNIEDGVFETLTNLELL (32)
>LRR6 206-227      LSLSFNSLSHVPPKLPSSLRKL (22)
>LRR7 227-251      LFLSNTQIKYISEEDFKGLINLTLL (25)
>LRR8 251-292      LDLSGNCPRCFNAPFPCVPCDGGASINIDRFAFQNLTQLRYL (42)
>LRR9 292-316      LNLSSTSLRKINAAWFKNMPHLKVL (25)
>LRR10 316-342     LDLEFNYLVGEIASGAFLTMLPRLEIL (27)
>LRR11 342-372     LDLSFNYIKGSYPQHINISRNFSKLLSLRAL (31)
>LRR12 372-399     LHLRGYVFQELREDDFQPLMQLPNLSTI (28)
>LRR13 399-423     INLGINFIKQIDFKLFQNFSNLEII (25)
>LRR14 423-486     IYLSENRISPLVKDTRQSYANSSSFQRHIRKRRSTDFEFDPHSNF YHFTRPLIKPQCAAYGKAL (64)
>LRR15 486-510     LDLSLNSIFFIGPNQFENLPDIACL (25)
>LRR16 510-535     LNLSANSNAQVLSGTEFSAIPHVKYL (26)
>LRR17 535-559     LDLTNNRLDFDNASALTELSDLEVL (25)
>LRR18 559-589     LDLSYNSHYFRIAGVTHHLEFIQNFTNLKVL (31)
>LRR19 589-613     LNLSHNNIYTLTDKYNLESKSLVEL (25)
>LRR20 613-644     LVFSGNRLDILWNDDDNRYISIFKGLKNLTRL (32)
>LRR21 644-669     LDLSLNRLKHIPNEAFLNLPASLTEL (26)
>LRR22 669-693     LHINDNMLKFFNWTLLQQFPRLELL (25)
>LRR23 693-717     LDLRGNKLLFLTDSLSDFTSSLRTL (25)
>LRR24 717-741     LLLSHNRISHLPSGFLSEVSSLKHL (25)
>LRR25 741-767     LDLSSNLLKTINKSALETKTTTKLSML (27)


TLR9 (manually measured from ladder positions)
>LRR1 68-92        LSLSSNRIHHLHDSDFAHLPSLRHL (25)
>LRR2 92-128       LNLKWNCPPVGLSPMHFPCHMTIEPSTFLAVPTLEEL (37)
>LRR3 128-148      LNLSYNNIMTVPALPKSLISL (21)
>LRR4 148-172      LSLSHTNILMLDSASLAGLHALRFL (25)
>LRR5 172-204      LFMDGNCYYKNPCRQALEVAPGALLGLGNLTHL (33)
>LRR6 204-225      LSLKYNNLTVVPRNLPSSLEYL (22)
>LRR7 225-249      LLLSYNRIVKLAPEDLANLTALRVL (25)
>LRR8 249-289      LDVGGNCRRCDHAPNPCMECPRHFPQLHPDTFSHLSRLEGL (41)
>LRR9 289-313      LVLKDSSLSWLNASWFRGLGNLRVL (25)
>LRR10 313-339     LDLSENFLYKCITKTKAFQGLTQLRKL (27)
>LRR11 339-369     LNLSFNYQKRVSFAHLSLAPSFGSLVALKEL (31)
>LRR12 369-396     LDMHGIFFRSLDETTLRPLARLPMLQTL (28)
>LRR13 396-420     LRLQMNFINQAQLGIFRAFPGLRYV (25)
>LRR14 420-477     VDLSDNRISGASELTATMGEADGGEKVWLQPGDLAPAPVD TPSSEDFRPNCSTLNFTL (58)
>LRR15 477-501     LDLSRNNLVTVQPEMFAQLSHLQCL (25)
>LRR16 501-526     LRLSHNCISQAVNGSQFLPLTGLQVL (26)
>LRR17 526-550     LDLSHNKLDLYHEHSFTELPRLEAL (25)
>LRR18 550-580     LDLSYNSQPFGMQGVGHNFSFVAHLRTLRHL (31)
>LRR19 580-603     LSLAHNNIHSQVSQQLCSTSLRAL (24)
>LRR20 603-633     LDFSGNALGHMWAEGDLYLHFFQGLSGLIWL (31)
>LRR21 633-658     LDLSQNRLHTLLPQTLRNLPKSLQVL

TLR10 (manually measured from ladder positions)
>LRR1 53-77        LDLSYNLLFQLQSSDFHSVSKLRVL (25)
>LRR2 77-101       LILCHNRIQQLDLKTFEFNKELRYL (25)
>LRR3 101-122      LDLSNNRLKSVTWYLLAGLRYL (22)
>LRR4 122-147      LDLSFNDFDTMPICEEAGNMSHLEIL (26)
>LRR5 147-170      LGLSGAKIQKSDFQKIAHLHLNTV (24)
>LRR6 170-193      VFLGFRTLPHYEEGSLPILNTTKL (24)
>LRR7 193-217      LHIVLPMDTNFWVLLRDGIKTSKIL (25)
>LRR8 217-247      LEMTNIDGKSQFVSYEMQRNLSLENAKTSVL (31)
>LRR9 247-274      LLLNKVDLLWDDLFLILQFVWHTSVEHF (28)
>LRR10 274-302     FQIRNVTFGGKAYLDHNSFDYSNTVMRTI (29)
>LRR11 302-331     IKLEHVHFRVFYIQQDKIYLLLTKMDIENL (30)
>LRR12 331-353     LTISNAQMPHMLFPNYPTKFQYL (23)
>LRR13 353-377     LNFANNILTDELFKRTIQLPHLKTL (25)
>LRR14 377-402     LILNGNKLETLSLVSCFANNTPLEHL (26)
>LRR15 402-426     LDLSQNLLQHKNDENCSWPETVVNM (25)
>LRR16 426-448     MNLSYNKLSDSVFRCLPKSIQIL (23)
>LRR17 448-471     LDLNNNQIQTVPKETIHLMALREL (24)
>LRR18 471-493     LNIAFNFLTDLPGCSHFSRLSVL (23)
>LRR19 493-517     LNIEMNFILSPSLDFVQSCQEVKTL (25)

Ribonuclease Inhibitor
>LRR1 28-56        VRLDDCGLTEEHCKDIGSALRANPSLTEL (29)
>LRR2 56-85        LCLRTNELGDAGVHLVLQGLQSPTCKIQKL (30)
>LRR3 85-113       LSLQNCSLTEAGCGVLPSTLRSLPTLREL (29)
>LRR4 113-142      LHLSDNPLGDAGLRLLCEGLLDPQCHLEKL (30)
>LRR5 142-170      LQLEYCRLTAASCEPLASVLRATRALKEL (29)
>LRR6 170-199      LTVSNNDIGEAGARVLGQGLADSACQLETL (30)
>LRR7 199-227      LRLENCGLTPANCKDLCGIVASQASLREL (29)
>LRR8 227-256      LDLGSNGLGDAGIAELCPGLLSPASRLKTL (30)
>LRR9 256-284      LWLWECDITASGCRDLCRVLQAKETLKEL (29)
>LRR10 284-313     LSLAGNKLGDEGARLLCESLLQPGCQLESL (30)
>LRR11 313-341     LWVKSCSLTAACCQHVSLMLTQNKHLLEL (29)
>LRR12 341-370     LQLSSNKLGDSGIQELCQALSQPGTTLRVL (30)
>LRR13 370-398     LCLGDCEVTNSGCSSLASLLLANRSLREL (29)
>LRR14 398-427     LDLSNNCVGDPGVLQLLGSLEQPGCALEQL (30)
```
